# Uip4 governs growth phase dependent organelle remodeling by modulating *Saccharomyces cerevisiae* lipidome

**DOI:** 10.1101/2023.03.31.535026

**Authors:** Pallavi Deolal, Kathirvel Ramalingam, Bhaskar Das, Krishnaveni Mishra

## Abstract

When yeast cells are exposed to nutrient-limiting conditions, they undergo transcriptional and translational reprogramming that results in the remodelling of metabolite utilization and organelle architecture. Organelle membranes and contacts also undergo structural and functional alterations. In the budding yeast Saccharomyces cerevisiae, regulated expression of Uip4 is shown to be a critical effector of nuclear shape and function, particularly during the stationary phase. In this work, we demonstrate that the absence of UIP4 affects the morphology of multiple other organelles including mitochondria, endoplasmic reticulum, vacuole and the distribution of lipid droplets. The results show that modulating carbon source, nitrogen availability and cellular energy state impact Uip4 expression. This expression of Uip4 is controlled by the transcription factor Msn2, downstream of Sch9 signaling pathway. Cells lacking Uip4 have poor survival in the stationary phase of the growth cycle. These cellular changes are concomitant with dysregulation of the global lipidome profile and aberrant organelle interaction. We propose that the dynamic and regulated expression of Uip4 is required to maintain lipid homeostasis and organelle architecture which is ultimately required to survive in nutrient limiting conditions such as stationary phase.

## Introduction

Lipid composition and concertation in a cell is dynamic and is altered during various stages of growth, growth in different culture media, calorie restriction and modified growth temperature (Klose et al., 2012; Mohammad et al., 2021). While most lipid classes are made in the cytoplasm and ER, mitochondria are also capable of synthesising phospholipids (Horvath and Daum, 2013). Processes involved in inter-organellar transport also play a critical role in the exchange of lipid species across membranes. Lipid transfer between organelles takes place via physical contact sites formed between various organelles (Gao and Yang, 2018; Helle et al., 2013; Henne, 2016; Cohen et al., 2018; Elbaz-Alon et al., 2014) and this directed transfer gives distinct lipid profiles to each organelle membrane and supports the functional specialization of organelles. Regulated expression and localization of enzymes involved in lipid metabolism is crucial for establishing membrane homeostasis (Mondal et al., 2022; Elbaz-Alon et al., 2015; Bahmanyar, 2015). Yeast and mammalian cells esterify sterols to form steryl esters, which are removed from membranes and packaged into lipid droplets (LDs). In addition to steryl esters, LDs also contain high levels of triacylglycerols (TG). These neutral lipids can be mobilized rapidly when cells need to form new membranes, for example, upon the resumption of growth. Lipid homeostasis is necessary for ensuring membrane integrity, the organization of the protein complexes on the membrane and hence functionality of membrane bound organelles. Organelle architecture changes are associated with perturbed concentration and distribution of various sub-classes of lipids (Wright et al., 1988; Hodge et al., 2010; Eiyama et al., 2013). While the pathways regulating fate of phosphatidic acid and membrane lipids are better understood, the regulation of inter-organellar lipid distribution and storage is not.

Budding yeast, *Saccharomyces cerevisiae* is a facultative anaerobe and in laboratory conditions, yeasts utilize glucose as the primary source of energy and grow exponentially while fermenting glucose that produces ethanol and acetate (Galdieri et al., 2010). Once the glucose in the medium is depleted, they undergo a short lag phase and begin to utilize ethanol and acetate from the medium released during the process of fermentation (Gray et al., 2004; Perez-Samper et al., 2018). After exhaustion of this carbon source, they enter a state of quiescence that is referred to as the stationary phase (Gray et al., 2004; Sun and Gresham, 2021). The cells undergo a considerable amount of change in total protein and transcript pool. Several proteins expressed largely during post diauxic shift and in stationary phase resemble those expressed under stress such as thermal, osmotic or nutrient (Gasch et al., 2000; Galdieri et al., 2010). Cell organelles also undergo structural and functional remodelling in response to the changed metabolic needs (Laporte et al., 2011).

In our recent work, we have shown that Uip4, an ER localized protein, is involved in regulation of nuclear morphology and is necessary to prevent clustering of NPCs (Deolal et al., 2022). The expression of Uip4 was higher in the cells at stationary phase which was concomitant with a decline in the nuclear pore complex quality and function. We took the investigation to study the role of Uip4 further by understanding its regulation and exploring other phenotypes. In this study, we report that Uip4 is a growth stage specific regulator of lipid homeostasis. We show that physiologically, expression of Uip4 is regulated by canonical nutrient signalling pathways of yeast and that the loss of *UIP4* leads to reduced survival in stationary phase. In addition, loss of *UIP4* affects mitochondrial morphology and function, and also results in lipid droplet accumulation. We find significant changes in concentration of various lipid classes in both logarithmic and post-mitotic phase of growth. Our results indicate that Uip4p plays a crucial role in nutrient driven organelle remodelling in *S. cerevisiae* by affecting cellular lipid profile and organelle contacts, thereby resulting in pleiotropic phenotypes. Future work will be aimed at unravelling the interactors and dissection the molecular mechanism.

## Results

### Uip4 expression in induced upon entry into stationary phase of growth

We observed that Uip4 expression in cells harvested from logarithmic and stationary phase varied (Deolal et al., 2022) and therefore wanted to assess the expression profile of Uip4 as the cell progresses along the various phases of growth. In order to understand the regulation of Uip4 expression, we harvested wild type cells having *UIP4*-13MYC tag and compared the level of protein expression at various phases of growth. We recorded the OD600 of the culture and plotted the growth curve to know the approximate growth phase (green series, right Y-axis Fig 1A). Corresponding to the marked sampling points, we measured the level of Uip4p in the biological replicates (Red series, left Y-axis Fig 1A). We found the Uip4p levels to be lowest during logarithmic phase when the cells are actively dividing and highest during the stationary phase when the growth is ceased (Fig 1A, Fig 1B and Deolal et. al. 2022). Our results reveal that Uip4 expression varies depending on the growth stage.

**Figure 1:**
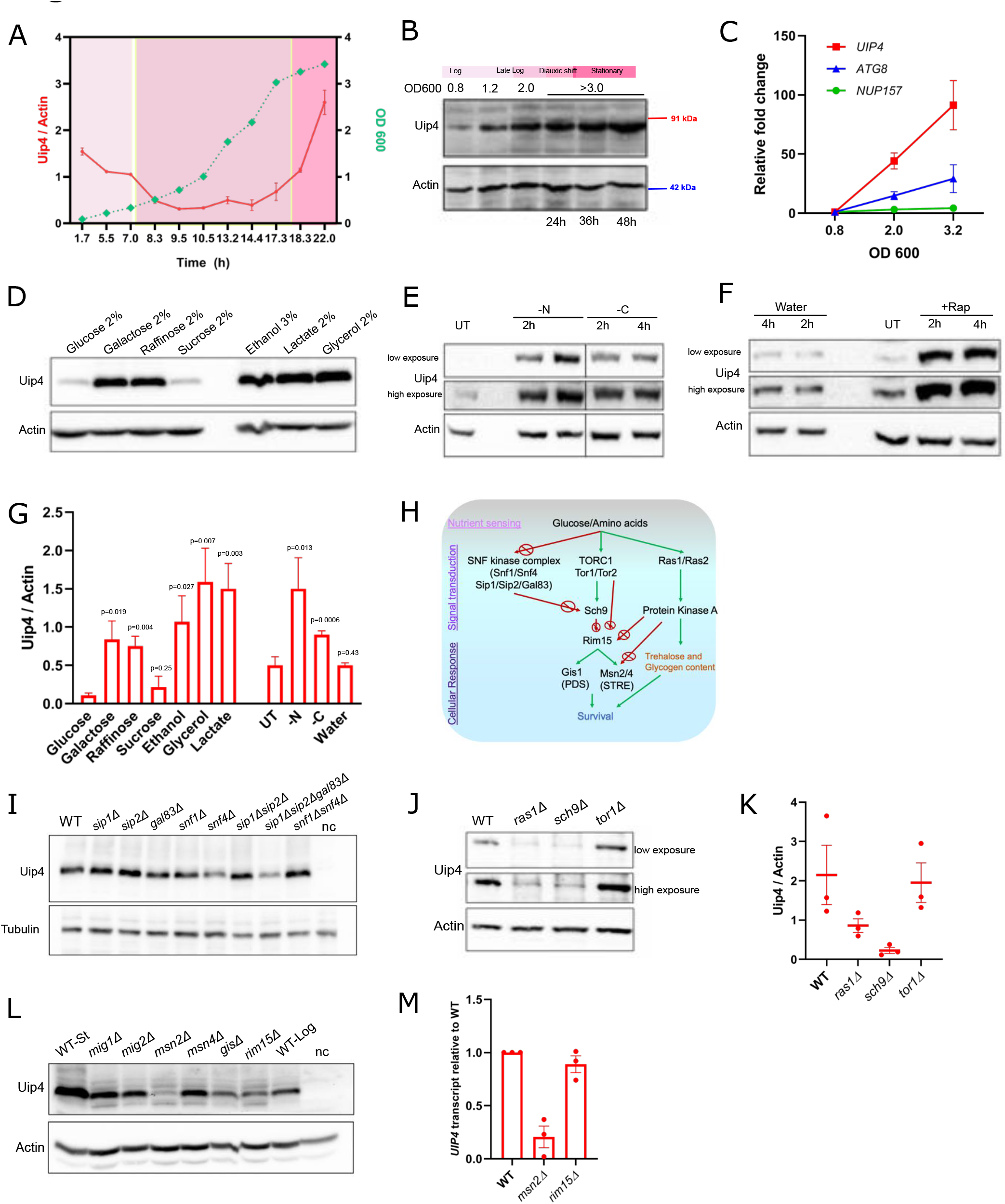
Uip4 expression is regulated in response to growth states. A. Wild type cells expressing Uip4-13xMyc from the endogenous loci were harvested at various time points from a liquid culture and Uip4 levels were checked by western blot using α-myc. α-actin was used as a loading control for total protein. The expression of Uip4 relative to actin was measured for each time point and an average from 3 independent experiment is plotted in left Y-axis (red curve). The corresponding OD600 of the harvested culture is plotted in the right Y-axis (green curve). B. A western blot representative of the expression of Uip4 from cells harvested at various phases of growth is shown. α-myc was used to detect Uip4 and α-actin was used as a loading control for total protein. Molecular weights are labelled on the right. C. RNA was isolated from WT cells at various stages of growth and qRT-PCR was performed using cDNA. The graph shows fold change in *UIP4* transcripts (red), *ATG8* (blue) and *NUP157* (green) relative to the level at mid-log phase (OD600 0.8). *ACT1* was used as normalizing control. n=3, error bars represent standard error of mean. D. Western blot showing the levels of Uip4 expression in the cells harvested from mid-log phase and transferred to a media containing indicated carbon source for 5 hours is presented. E, F. Wild type cells expressing Uip4-13xMyc were grown to mid-log phase and the culture was split into various sets for further treatments as indicated. α-myc was used to detect Uip4 and α-actin was used as a loading control for total protein. G. The bar graph shows quantification of Uip4 expression relative to actin from the growth conditions described in D-F. n=3, error bars represent standard error of mean. H. The schematic shows nutrient sensing pathways in yeast. I, J. Indicated strains expressing Uip4-13xMyc from a centromeric plasmid under the control of its native promoter were harvested and Uip4 levels were checked by western blot. α-myc was used to detect Uip4 and α-actin was used as a loading control for total protein. K. The bar graph shows quantification of Uip4 expression relative to Actin from the strains described in I. n=3, error bars represent standard error of mean. L. Indicated strains expressing Uip4-13xMyc from a centromeric plasmid under the control of its native promoter were harvested and Uip4 levels were checked by western blot. α-myc was used to detect Uip4 and α-actin was used as a loading control for total protein. WT-Log and WT-St are extracts from wild type cells harvested either at mid-log phase or stationary phase of growth. nc is for negative control which is extract from wild type cells transformed with empty vector. M. RNA was isolated from indicated strains and qRT-PCR was performed using cDNA. *ACT1* was used as normalizing control. The histogram shows abundance of *UIP4* transcripts in *msn2*Δ and *rim15*Δ relative to WT.

To test if this regulation was transcriptionally controlled, we quantified the abundance of *UIP4* transcripts by quantitative RT-PCR (Fig 1C). As controls, we tested the transcriptional status of *ATG8,* an autophagy related protein and *NUP157,* a nucleoporin. Autophagy is induced upon glucose depletion and entry into stationary phase while the expression of nucleoporins remains fairly unchanged (Galdieri et al., 2010; Toyama et al., 2013). As anticipated, we find that levels of *NUP157* mRNA do not alter significantly between logarithmic (OD600-0.8) to stationary phase (OD 600-3.0) while transcription of *ATG8* is induced (Fig 1C). *UIP4* mRNA increases several folds as the cells exit exponential growth phase and enter stationary phase. This suggests that expression of Uip4 is transcriptionally controlled.

Yeast utilizes glucose as the preferred carbon source and grows logarithmically while metabolizing glucose to ethanol (Perez-Samper et al., 2018). Presence of glucose in the growth medium affects various cellular processes such as expression of genes involved in mitochondrial respiration, gluconeogenesis and lifespan regulation (Rolland et al., 2002; Perez-Samper et al., 2018). One of the key changes that occurs during transition from logarithmic to stationary phase is the shift from fermentation to respiration (Galdieri et al., 2010). Genes repressed under glucose abundance are also involved in utilization of alternate carbon sources (Santangelo, 2006). In order to test if Uip4 is involved in this metabolic shift or responsiveness to altered metabolic demand, we analysed Uip4 levels in cells grown in alternate carbon sources. Uip4 expression was induced upon shift to both fermentable and non-fermentable carbon sources, except for sucrose (Fig 1D, 1G). In budding yeast, the monosaccharides glucose and fructose generated upon hydrolysis of sucrose enter the glycolytic pathway. Therefore, glucose and sucrose result in repression of mitochondrial function. During sucrose metabolism, the glucose repression is more effective compared to other alternate carbon sources (Santangelo, 2006). Therefore, it appears that the induction of Uip4 is concomitant with decline in glycolytic repression and dependence on mitochondrial function.

### Uip4 expression in induced upon growth in nutrient limiting conditions

Nutrients become limiting in stationary phase. Having established that Uip4 expression is upregulated in stationary phase and media containing alternate carbon sources, we directly depleted either carbon or nitrogen source in the growth media and looked at Uip4 levels. We subjected cells harvested from logarithmic phase grown in 2% glucose containing media to either carbon (C) starvation (0.75%SC, 0.05% glucose) or nitrogen (N) starvation (0.17%YNB, 2% Glucose). Either C or N starvation induced Uip4 expression (Fig 1E, Fig S1A). A greater induction was observed upon N starvation as compared to C starvation. Rapamycin treatment also results in starvation like phenotype, so we assessed Uip4 expression upon treatment with rapamycin. Addition of rapamycin resulted in an even higher induction of Uip4 expression (Fig 1F, Fig 1G, Fig S1B). Interestingly, the protein levels remain unchanged when cells are left in water (Fig 1F, 1G). This indicates that a specific signalling response induced by nutrient sensing regulates Uip4 expression.

### Induction of Uip4 expression is regulated by the RAS/PKA and Sch9 kinase signalling

To understand how the nutrient levels are communicated for Uip4 expression, we studied the protein level of Uip4 in various single gene deletion mutants of key kinases that are implicated in nutrient sensing in yeast. Three main pathways involving the protein kinases PKA, TOR and Sch9 integrate gene expression with the nutrient sensing cues from the growth medium in a Rim15 dependent manner (Toda et al., 1988; Rolland et al., 2002; Swinnen et al., 2006) (Fig 1H). Rim15 is a glucose-sensing protein kinase required for entry of cells into stationary phase (Reinders et al., 1998). Activation of Rim15 under nutrient depletion and stress conditions regulates growth and controls gene expression in a Msn2/4 and Gis1 transcription factor dependent manner (Martínez-Pastor et al., 1996; Pedruzzi, 2000). Additionally, Snf1/Snf4 kinase complex (AMP-activated protein kinase of mammals) is involved in expression of various glucose-repressible genes (Jiang and Carlson, 1997; Hedbacker and Carlson, 2008). Yeast Snf1 is activated during low nutrient availability and energy limiting conditions, thereby promoting catabolic activities. The Snf1 kinase complex of yeast includes the activating subunit, Snf4, and the bridging proteins Sip1, Sip2 and Gal83 (Jiang and Carlson, 1997). Sip1, Sip2 and Gal83 are essential for regulating interaction of Snf1/4 kinase complex with specific substrates (Yang et al., 1994). As the cells shift from glucose dependent fermentation to either an acetate or ethanol utilising respiration during the stationary phase, several pathways centred around Snf1 kinase are influenced. This includes expression and nucleocytoplasmic translocation of transcription factors involved in expression of genes required for utilization of alternate carbon sources (Hedbacker and Carlson, 2008).

We tested which of these canonical pathways influence the regulation of expression of Uip4. We transformed the deletion mutants with centromeric plasmid expressing *UIP4* tagged with a *13MYC* epitope at its C terminal, under its endogenous promoter. The expression of Uip4 from the plasmid was similar to the expression from a strain bearing tag at genomic locus (Fig S1C). We find that Uip4p expression is attenuated in *sip1*Δ*sip2*Δ*gal83*Δ and *snf4*Δ cells only, and not in either of the single deletion mutants (Fig 1I). In the absence of either Ras1, a GTPase involved in activation of PK-A, or Sch9 kinase, Uip4 levels are low as compared to the wild type (Fig 1J). While in *ras1*Δ, we observe an increase in Uip4 expression during stationary phase as compared to the levels in logarithmic phase, such an induction is not seen in case of *sch9*Δ (Fig S1D). This suggests that Sch9 is one of the key upstream kinases that partakes in regulating Uip4 expression as the cells transition to stationary phase. Activation of Tor1/Tor2 kinase complex takes place during nutrient abundant conditions, thereby promoting anabolic processes. Our results suggest that TOR mediated signalling is dispensable for Uip4 expression as we do not see any major reduction or upregulation in Uip4 expression in *tor1*Δ when compared to wild type (Fig 1J, 1K). Similar signalling pathways also seem to regulate the abundance of Uip4 during either glucose (Fig S1E, S1F) or nitrogen depletion (Fig S1G).

To identify the potential transcription factor that activates the expression of Uip4p, we analysed the promoter of *UIP4* for transcription factor binding sites using the Promoter Analysis tool of Yeastract (Monteiro et al., 2020). We found various possible binding sites for transcription factors Msn2 and Gis1. Both of these transcription factors are implicated in upregulation of stress responsive genes in stationary phase of yeast growth (Pedruzzi, 2000; Durchschlag et al., 2004). Apart from Msn2 and Gis1, we also measured Uip4 expression in the absence of Msn4, Mig1 and Mig2. Msn4 is also a stress responsive transcription factor, with several overlapping targets with Msn2 (Martínez-Pastor et al., 1996). Mig1 and Mig2 are involved in control of glucose-repressible genes. We find maximum downregulation of Uip4p in *msn2*Δ as compared to either *msn4*Δ or *gis1*Δ (Fig 1L). We do not find any effect of either *mig1* or *mig2* deletion on the expression of Uip4. We confirmed the downregulation in *msn2*Δ at the transcript level by performing quantitative RT-PCR. The abundance of *UIP4* mRNA in *msn2*Δ is highly reduced as compared to the wild type cells (Fig 1M). Since the activity of Msn2 is largely dependent on the upstream kinase Rim15, we asked if the downregulation is mediated via Rim15. However, we find only a modest reduction in the Uip4 expression in *rim15*Δ (Fig 1L). Likewise, the transcript levels also do not change much as compared to the wild type (Fig 1M). This hints towards Msn2 mediated Uip4 regulation by upstream regulators other than Rim15.

### Uip4 is essential for viability of non-dividing yeast cells

While loss of Uip4 does not affect the mitotic growth rate of cells when grown in glucose containing media (Deolal et al., 2022), we tested growth in alternate carbon sources which affect the expression of Uip4. We sub-cultured WT and *uip4*Δ cells to media containing different carbon sources and monitored the growth profiles. We did not find any major difference in the mitotic growth profiles of the two strains in fermentable (Fig S2A) and non-fermentable carbon source containing medium. In order to test the significance of Uip4 induction in post-mitotic/ stationary phase, we looked at the survival of wild type and *uip4*Δ cells in the post-mitotic state. To test the ability to survive long term without nutrient replenishment, CFU/ml (Colony forming units) for each strain was quantified and normalised to the number of survivors for a 1-day old culture. The fraction of survivors was calculated for each strain and we find a noticeable reduction in the viability of *uip4*Δ cells compared to wild type as the number of days increase (Fig 2A). Some of the cellular changes that occur during stationary phase overlap with those induced during stress such as heat shock (Li et al., 2013). In order to test if Uip4 has any protective effect at increased temperature as well, we performed growth assays on agar plate. Equal number of wild type and *uip4*Δ cells were spotted on SC plate and then grown at either 30°C or 39°C as well. We find that loss of *UIP4* had a negative impact on the growth at 39°C (Fig 2B, Fig S2C). We also noticed that the *uip4*Δ cells in stationary phase arrest growth with small sized buds as compared to the predominantly unbudded wild type cells (data not shown). There was no significant difference in the overall cell size between WT and *uip4*Δ cells in the early stages of growth but in the later stages of growth, cells lacking *UIP4* were significantly larger than wild type (Fig S2D). Taken together, these results suggest that elevated expression of Uip4 in late stages of growth is important for cell survival in the post-mitotic phase.

**Figure 2:**
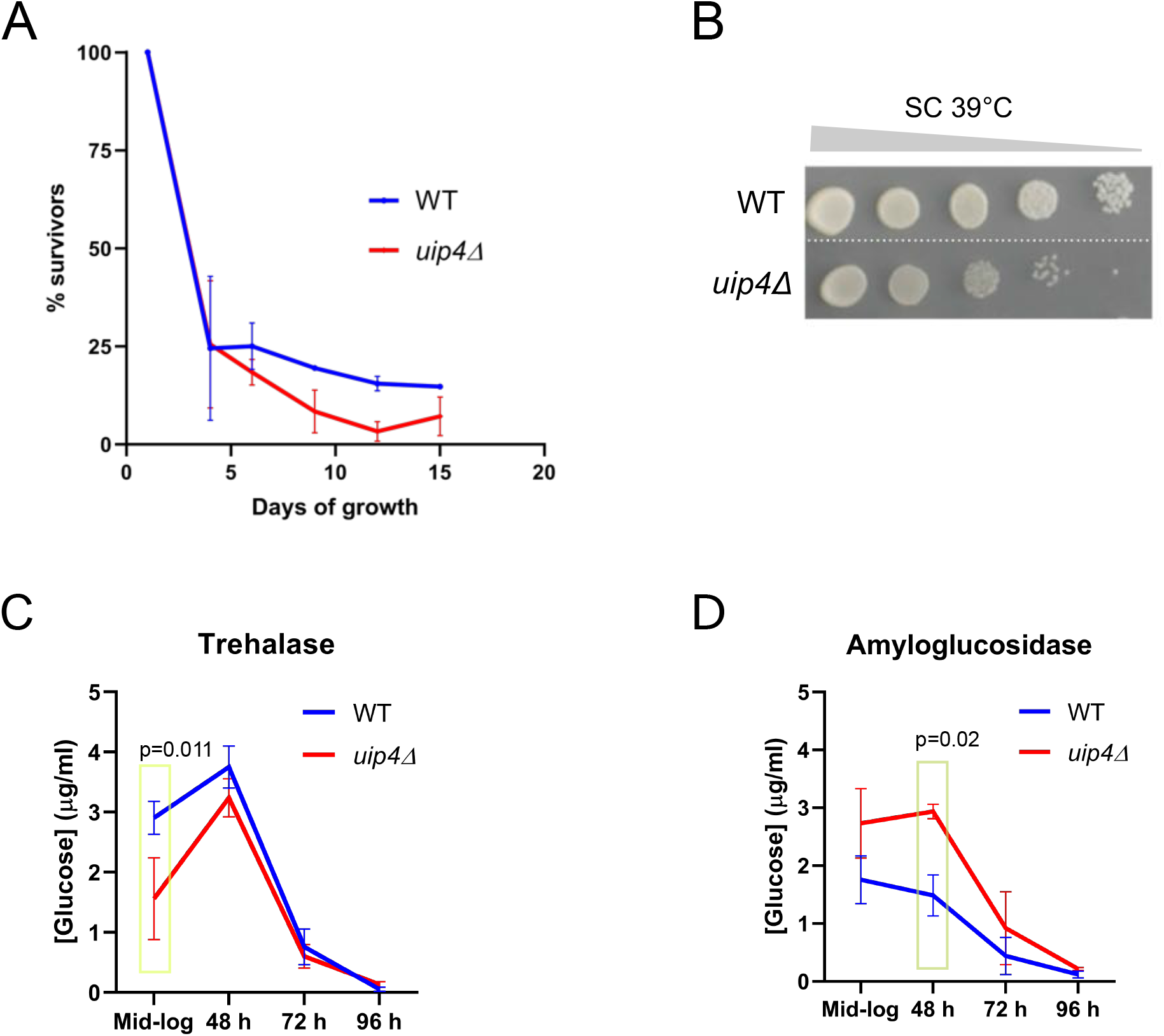
A. The fraction of cells surviving in WT and *uip4*Δ as the time progresses is plotted. The survivors on day 1 after subculture was considered as 100%. B. Serially diluted cultures of the indicated strains were taken and 5µl of 10-fold serial dilutions were spotted on a SC-plate. The plates were incubated at either 30°C for 2 days or 39°C for 3 days. C,D. Cellular storage carbohydrates trehalose and glycogen were estimated by measuring release of glucose by enzymatic digestion with trehalase and amyloglucosidase respectively, in WT and *uip4*Δ at indicated stages of growth.

Two of the key carbohydrate energy reserves in yeast are glycogen and trehalose (Parrou et al., 1997). Trehalose, a non-reducing disaccharide, has a role in attenuating the proteotoxicity that occurs during stationary phase and promoting stress tolerance. This protective role of trehalose is known to promote longevity of yeast cells (Samokhvalov et al., 2004; Kyryakov et al., 2012). While the glycogen accumulation peaks early at diauxic shift, maximal trehalose accumulation occurs in the stationary phase (Li et al 2013). We tested whether there are differences in their utilization and the total stored levels of these storage carbohydrates which could lead to reduced survival in stationary phase of *uip4*Δ. Despite lower trehalose content in the log-phase, accumulation of trehalose in the stationary phase was not affected in *uip4*Δ cells (Fig 2C). On the other hand, there was a slight, yet significant, increase in the glycogen reserves of *uip4*Δ cells (Fig 2E). The increase in size of *uip4*Δ cells noted in stationary phase could be a consequence of this increased glycogen accumulation (Fig S2D, Fig 2D). However, these differences were only modest and the utilization of the reserve carbohydrates also did not seem to be impaired by loss of *UIP4.*

### *Uip4*Δ cells have abnormal organelle morphology

Apart from metabolic changes, several architectural and organizational changes also take place as the cells transition from logarithmic phase of growth to stationary phase (Li et al., 2013; Picard et al., 2013; Galloway et al., 2012; Okamoto and Shaw, 2005; Kutay et al., 2014). Architectural changes to organelles are a part of nutrient sensing response (Sun and Gresham, 2021; Sagot and Laporte, 2019). Nuclear size and circularity, are also reported to change when carbon source is altered or when cells are shifted to starvation inducing media (Wang et al., 2016). Mitochondria display a highly dynamic and heterogenous morphology in a population of cells and, the shape and number of mitochondria in a cell are indicative of the energy states and respond to genetic and environmental perturbations (Shutt and McBride, 2013) (Wai and Langer, 2016). ER has an extensive network of sheets and tubules. Unique and dynamic contact sites formed between ER and other organelles are important for lipid transfer, exchange of Ca^2+^ and inter-compartment shuttling of metabolites (Zhang and Hu, 2016). Maintenance of proper ER morphology is an important determinant in establishing membrane contacts and thereby enabling organellar functions (Zhang and Hu, 2016; Cohen et al., 2018). The lipid composition also changes in response to metabolic needs and stress adaptation (Li et al., 2013; Casanovas et al., 2015). Neutral lipids are important for survival in stationary phase (Hariri et al., 2018).

We have previously shown that Uip4 localizes to NE/ER (Fig S3A) and the localization of Uip4 does not change between logarithmic and stationary phase (Deolal et al., 2022). Since Uip4 is localized to ER which is in continuity with the nuclear envelope, we first tested if ER morphology is altered in *uip4*Δ. Clearly, in the absence of Uip4 we observed disruption of ER structure (Fig 3A, 3B). The luminal ER marker dsRed-HDEL is distributed to nuclear, peripheral and cytosolic ER in wild type and appears as two prominent rings in addition to some cytosolic ER (Fig 3A, WT). On the other hand, we observed disrupted cortical ER and abnormal perinuclear accumulation of the luminal marker in *uip4*Δ (Fig 3A, *uip4*Δ*-*yellow arrows). To further confirm the membrane deformations, we used a GFP tagged integral ER membrane protein Scs2, known to interact with tethering factors and formation of cortical ER (Loewen et al., 2007; Chen et al., 2012). We found that in the absence of Uip4p, the cortical ER is less continuous and regions lacking Scs2 could be seen (Fig 3B, white arrows). Excess membrane accumulation can be observed in regions adjacent to nucleus and regions connecting nuclear and cortical ER (Fig 3B, yellow arrows). This suggests that the balance of ER membrane distribution at the nuclear and cortical regions is disrupted in the absence of Uip4.

**Figure 3:**
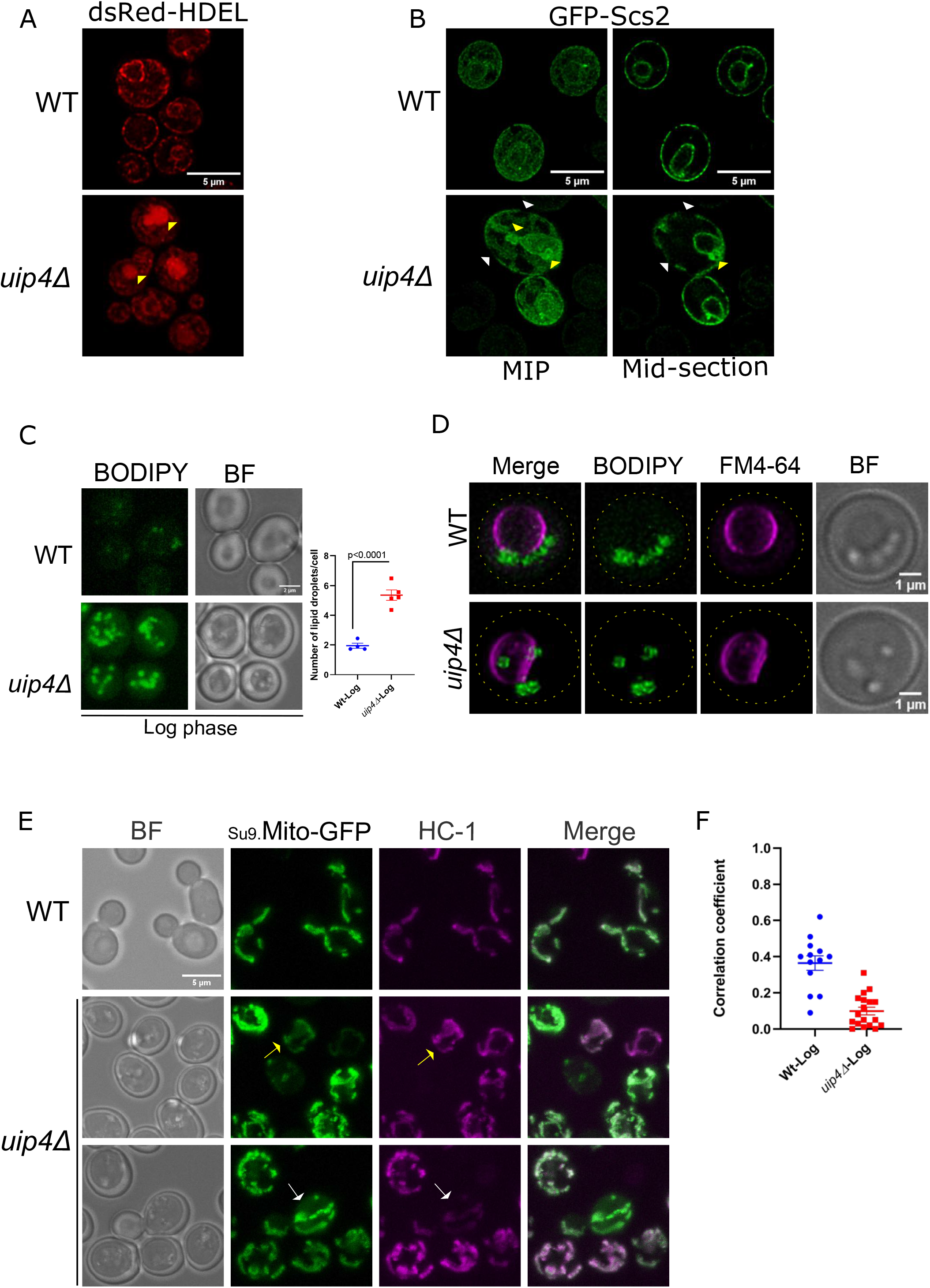
A, B. ER morphology was assessed in wild type and *uip4*Δ cells expressing either dsRed-HDEL or GFP-Scs2 during mid-log phase of growth. Cells representative of the ER morphology displayed by maximum fraction of cells is shown. Scale-5µm. MIP-maximum intensity projection. C. Lipid droplets in WT and *uip4*Δ harvested at mid-log phase of growth were stained using BODIPY 493/503. Maximum intensity projections of representative images are shown. Scale-2µm. A plot showing distribution of number of lipid droplets per cell is shown on the right. Horizontal line indicates mean. D. WT and *uip4*Δ cells were co-stained with BODIPY and FM4-64 to visualize lipid droplets and vacuole membrane respectively. Deconvolved images representative of the distribution are shown and the scale bar is labelled. Cell outline in the merged panel is shown in yellow. E. Mitochondrial morphology was assessed in wild type and *uip4*Δ cells harbouring a plasmid expressing Su9-Mito-GFP harvested during mid-log phase of growth. Cells were additionally stained with a potentiometric mitochondrial dye HC-1. Representative images are shown. Arrows mark cells that have differential staining of the mitochondria targeted protein (Su9-Mito-GFP) and the dye (HC-1). Scale -5µm. Cells marked with white and yellow arrows are indicative of heterogenous staining. F. Degree of colocalization between mitochondria targeted protein (Su9-Mito-GFP) and the dye (HC-1) was estimated and the values are plotted. Horizontal line represents median.

Another organelle in yeast associated with ER is the lipid droplets that bud off from ER and acts as a channel for lipid transfer between organelles. Therefore, intuitively we also looked at the number and spatial distribution of lipid droplets in wild type and *uip4*Δ cells. During the logarithmic growth phase, where the lipids are being utilized for membrane biosynthesis to support cell growth, in the wild type cells, lipid droplets are few and small in size (Fig 3C). When the cells are in stationary phase of growth characterized by growth cessation, lipid storage is preferred over membrane biosynthesis (Czabany et al., 2006). This results in an increase in the number of lipid droplets in wild type (Fig S3B, Fig 3D). In the stationary phase, most of these lipid droplets in WT cells were organized as a string along the vacuole membrane (Fig 3D). Occasionally, more than one such clusters were observed in addition to a few cytoplasmic lipid droplet clusters (Fig S3B, Fig 3D). Irrespective of the distribution, the lipid droplets remained spatially restricted in wild type. We found significant differences in the lipid droplet number and distribution in the absence of *UIP4* (Fig 3C, 3D). Cells lacking *UIP4* had multiple, large lipid droplets when compared to wild type cells in both logarithmic phase as well as in stationary phase (Fig 3C, Fig S3B). In addition, the lipid droplets were highly mobile and irregularly distributed *uip4*Δ cells (Fig 3D). A small fraction of lipid droplets were also found to be encapsulated within the folds of vacuole membrane in *uip4*Δ. These data show that there is a significant alteration of both number and organization of lipid droplets.

### Loss of *UIP4* leads to abnormal mitochondrial morphology and function

Mitochondrial function is largely dependent on its structure and membrane potential (Wai and Langer, 2016) which is responsive to cellular cues such as nutrient starvation and oxidative stress (Wai and Langer, 2016; Shutt and McBride, 2013). Respiration-linked genes are specifically upregulated in the stationary phase of growth (Kaniak-Golik and Skoneczna, 2015). Since Uip4 is expressed during respiration requiring conditions and we see various membrane related changes in NE, ER, vacuole and lipid droplets, we sought to investigate if there are differences in mitochondrial membrane morphology and/or membrane potential.

In order to examine the mitochondrial morphology we used ectopically expressed Su9-Mito-GFP which is imported to the mitochondrial matrix in a mitochondrial membrane potential dependent manner (Rapaport et al., 1998). In addition to this we assessed the direct uptake of a mitochondrial membrane potential dependent dye-HC1(Raja et al., 2021, 2017). In a wild type cell, the mitochondria appears like a continuous filamentous structure while in the absence of *UIP4*, mitochondria was found to be either fragmented or clustered (Fig 3E). There was no significant difference in the overall signal intensity for either protein or dye retained in the mitochondria of wild type and *uip4*Δ cells (Fig S3C). In wild type, the dye signal overlapped almost entirely with the Su9-Mito-GFP in all the cells. *Uip4*Δ on the other hand, displayed more variability in the dye uptake (Fig 3E, yellow and white arrows), resulting in less correlation between the GFP and dye staining (Fig 3F). This hints at the functional heterogeneity in the population of mitochondria in *uip4*Δ cells.

Mitochondria plays an important role in orchestration of the metabolic reprograming as the cells transition through the diauxic shift which involves morphological remodeling as well as shift from fermentation to respiration (Okamoto and Shaw, 2005; Shutt and McBride, 2013). In order to understand the dynamic regulation of mitochondrial morphology, we followed the morphological differences in wild type and *uip4*Δ cells through the growth stages. We performed live cell imaging of cells harvested at multiple stages of growth as marked in Fig 4A and the distribution of these distinct populations is represented in the graph shown in Fig 4B. Loss of tubular structure was seen in wild type cells as they progressed towards later stages of stationary phase of the growth cycle (Fig 4B). We observed that in agreement with previous reports the mitochondria move towards the cellular periphery in later stages of growth (Fig 4E-48h, 96h). In early phases of growth of *uip4*Δ, mitochondria lose their tubular architecture and are fragmented. There is also a population of mitochondria that is interconnected and dense (Fig 4A, B). In post-mitotic phase, unlike in wild type cells, in *uip4*Δ, fragmented mitochondria do not move to the periphery and importantly, about 15% of the cells in the population retain networked mitochondria.

**Figure 4:**
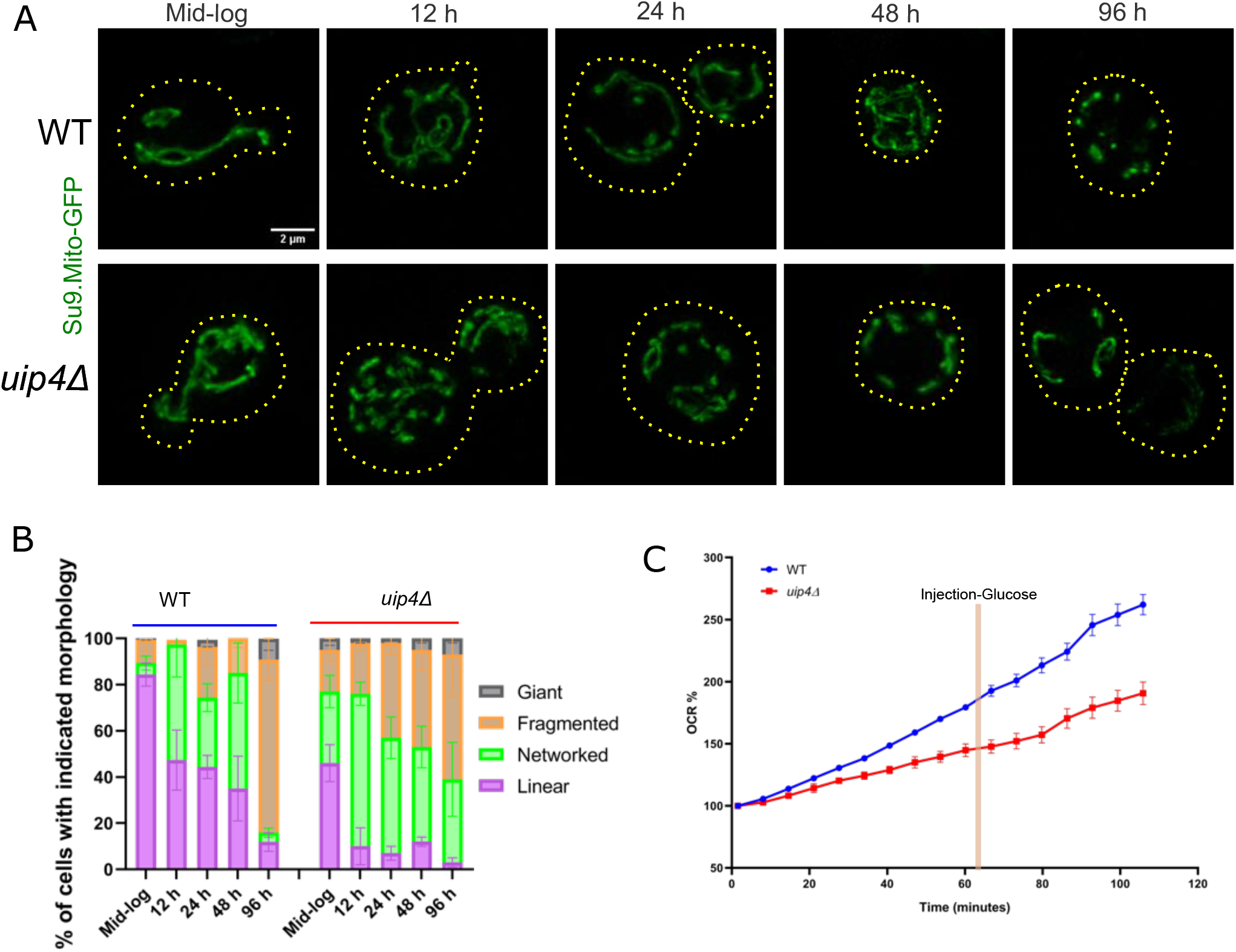
A. Mitochondrial morphology was assessed in wild type and *uip4*Δ cells harbouring a plasmid expressing Su9-Mito-GFP during various phases of growth as indicated on the top of each image panel. Cells representative of the mitochondrial morphology displayed by maximum fraction of cells is shown. Scale-2µm. B. The various morphologies of mitochondria present in the wild type and *uip4*Δ were broadly classified in to four categories and a distribution of these is plotted in the stacked bar graph. C. The mitochondrial respiration of wild type and *uip4*Δ cells was tested by directly measuring the oxygen consumption rate (OCR) of the cells on the Seahorse XFp Analyzer. The ECAR and OCR profiles normalized with values at the beginning of the workflow as baseline are plotted. The injection is an addition of glucose (111mM) in the medium.

We set out to test if there are any differences in the fermentative and respiratory output between wild type and *uip4*Δ cells due to morphological differences. We used the Seahorse XF technology to obtain measurements of extracellular acidification rate (ECAR) and oxygen consumption rate (OCR), simultaneously, in live cells under basal conditions as a measure of the metabolic state. Glucose in the medium is converted to lactate during glycolysis and this results in acidification of the assay medium. We did not observe any major difference in the ECAR between wild type and *uip4*Δ cells (Fig S3D). On the contrary, the OCR of *uip4*Δ cells was lower than the wild type cells (Fig 4C). This reduction in mitochondrial respiratory capacity hints at impaired mitochondrial function in the absence of *UIP4*. These data show that in the absence of Uip4, both the structure and functional competence of mitochondria are compromised.

### Organelle contacts are dysregulated in the absence of Uip4

Inter-organellar contacts play important roles in exchange of metabolites including lipids between organelles, and multiple such constitutive and regulated contact sites have been observed. Contact sites formed by association of lipid droplets are essential in metabolic rewiring (Radulovic et al., 2013; Cohen et al., 2018; Dakik and Titorenko, 2016). We investigated the sub-cellular distribution of lipid droplets in relation to mitochondria and vacuole during the logarithmic phase of growth since *uip4*Δ cells had abnormally high lipid droplets. In wild type, the lipid droplets were not associated either with mitochondrial or vacuole membrane (Fig 5A, S3E). Thus, during log-phase of growth the contacts between mitochondria, vacuole and lipid droplets are limited in wild type cells. On the other hand, the abnormally increased lipid droplets in *uip4*Δ show increased association with vacuole and mitochondria (Fig 5A, S3F). Some of the mitochondrial signal is also found in vacuole proximal regions (Fig S3F, grey arrow). The structural changes were also reiterated in 2D transmission electron microscopy. Lipid droplets were visible in very few sections of the wild type electron micrographs (Fig 5B, left). In contrast, the lipid droplets were very prominent in the *uip4*Δ electron micrographs (Fig 5B, right). These lipid droplets were primarily collocated at the nuclear-vacuole junctions and between vacuole membrane whorls.

**Figure 5:**
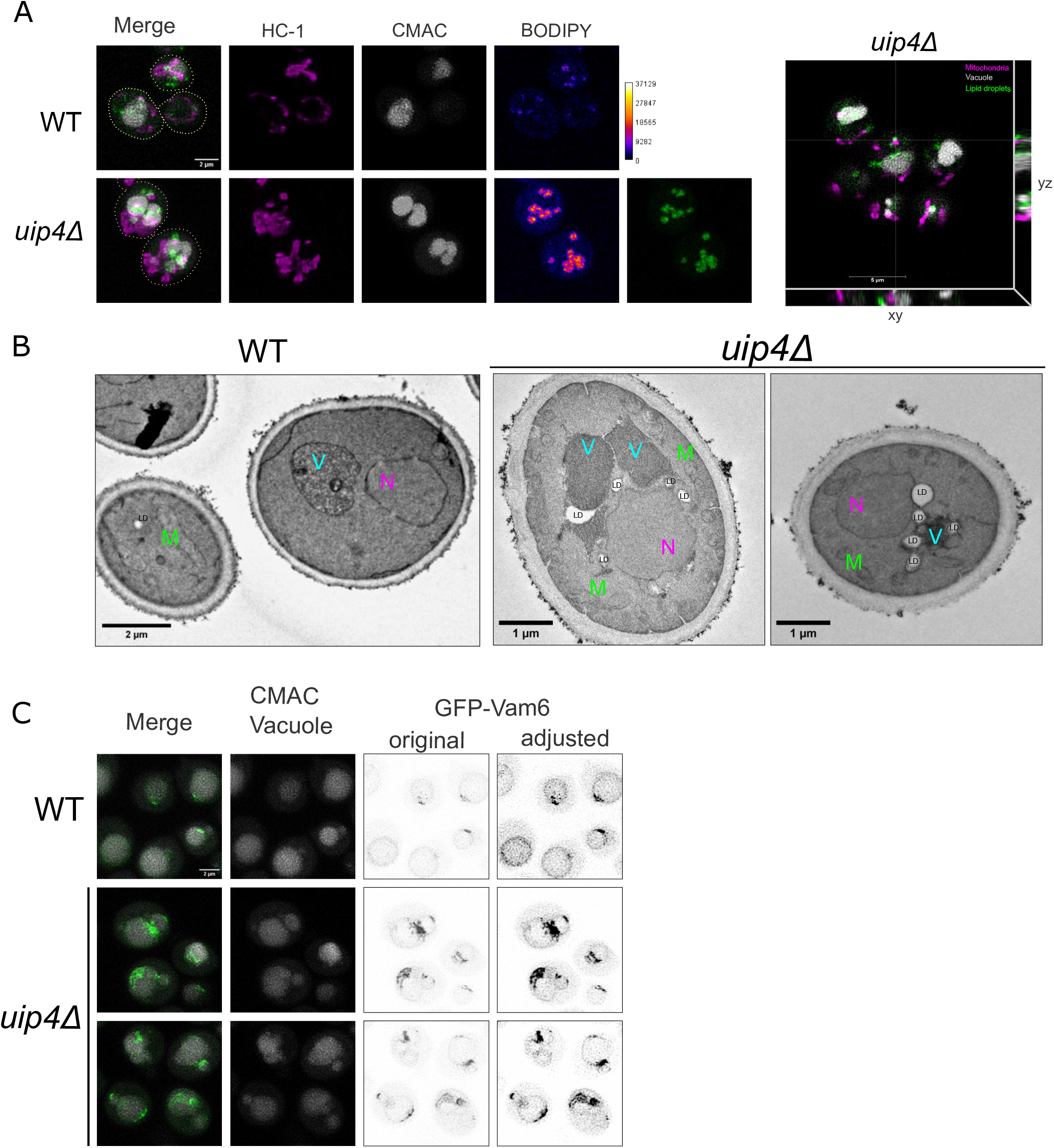
A. WT and *uip4*Δ cells harvested during logarithmic phase of growth were co-stained with HC-1, BODIPY and CMAC-blue to label mitochondria, lipid droplets and vacuole lumen respectively. The relative distribution of the three signals as seen in majority of the cells is shown in the panels. Scale-2µm. Cells are outlined in yellow dotted lines. Fire LUT is used for BODIPY to visualize the signal better. The intensity scale is shown adjacent to it. An orthogonal section showing the overlap of the signals for *uip4*Δ is shown on the right. B. TEM was performed for WT and *uip4*Δ cells. 2D-electron micrographs for WT and *uip4*Δ cells representative of the organelle (nucleus-N, mitochondria-M, vacuole-V and lipid droplets-LD) morphology are shown. Scale bars are labelled on the images. C. WT and *uip4*Δ cells expressing GFP-Vam6 harvested during logarithmic phase of growth were co-stained with CMAC-blue to label vacuole lumen. A black and white inverted LUT is used for GFP-Vam6 to better represent the differences in the overall signal and distribution in WT and *uip4*Δ. Scale-2µm

Contacts between lipid droplets and vacuole membrane are important in regulating lipid homeostasis and sterol metabolism, particularly during the stationary phase (Cohen et al., 2018; Henne, 2016; Dakik and Titorenko, 2016). Lipid droplets act as sites of lipid and protein exchange as an adaptation mechanism during metabolic rewiring (Radulovic et al., 2013; van Zutphen et al., 2014). Vam6 (also referred as Vps39), is an essential component of the vacuole and mitochondrial contact site in yeast (Elbaz-Alon et al., 2014). Vam6 is required for formation of the vacuole and mitochondria patch (vCLAMP). These contact sites are regulated in response to cellular metabolic state (Hönscher et al., 2014). vCLAMPs are believed to be important during growth in glucose containing media and their prominence is reduced during increased respiratory demands (Hönscher et al., 2014). Since we observed increased apposition of mitochondria and lipid droplets with vacuole, we asked if the vacuole -mitochondria contact sites are also extended by visualizing the Vam6 distribution. In logarithmically growing wild type cells, Vam6 marks the vacuole membrane and is enriched at one or two sites where vCLAMPs are formed (Fig 5C, Fig S3G). In the absence of *UIP4*, we find that these sites are enlarged (Fig 5C) and the degree of overlap between mitochondrial and vacuole membrane is also more as compared to wild type (Fig 5C, S3H). Altogether, it shows that there is abnormal accumulation and distribution of lipid droplet within subcellular spaces, with increased proximity to other organelles, in the absence of Uip4.

### Loss of Uip4 causes cellular lipid imbalance

We reasoned that the increased lipid droplets and altered morphologies of organelles could be the result of impaired lipid homeostasis. To this end, we assessed the effect of loss of *UIP4* on the global lipidome profile of yeast during mid-log as wells as in stationary phase of growth. When compared to the mammalian system, the yeast lipidome is relatively simple (Klose et al., 2012). However, there is huge diversity and flexibility in the various classes of lipids in yeast as well. There is a large variability in the amount and kind of lipids distributed within the sub-cellular spaces (Klose et al., 2012). As the yeast cells progress from logarithmic to stationary phase, notable changes of the lipidome take place (Fig 6A) typically leading to increase in triacylglycerol synthesis and lipid droplet formation (Klose et al., 2012; Wang, 2014; Barbosa et al., 2019).

**Figure 6:**
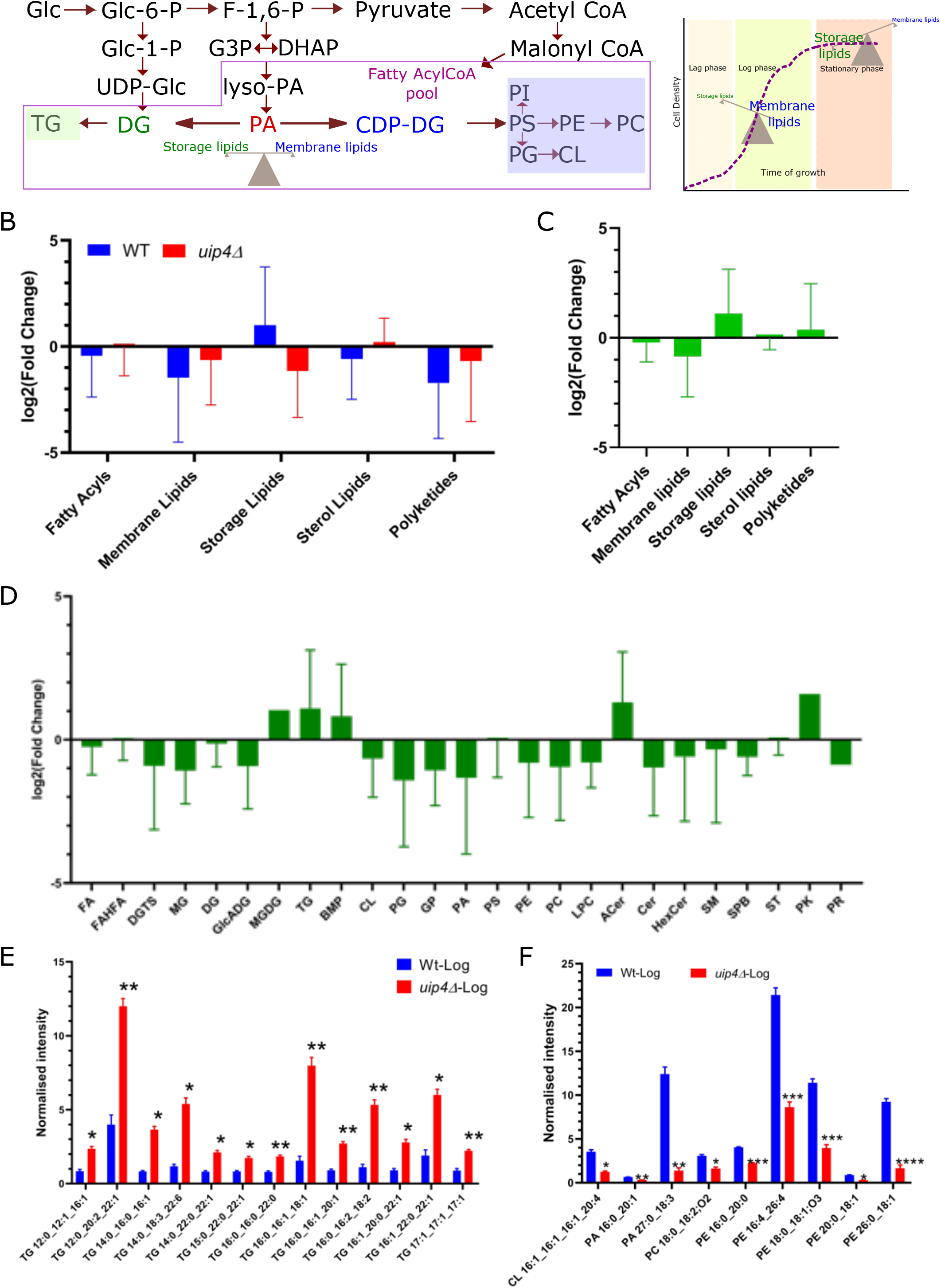
A. The schematic shows a simplified representation of the synthesis of storage and membrane lipid in yeast starting from the primary energy source-glucose. The fatty acylCoA pool is a primary substrate for most pathways that use fatty acids either for synthesis of complex lipids or for generation of energy. Phosphatidic acid (PA) levels are highly regulated by various enzymes and specific sub-cellular distribution. Conversion of PA to diacyl glycerol (DG) is an important regulatory step in the biosynthesis of storage lipid-triacylglycerol (TG) and other membrane phospholipids like phosphatidylserine (PS), phosphatidylcholine (PC), cardiolipin (CL) etc. The graph on the right shows the shift in the dynamics of storage versus membrane lipids as the yeast cells transition from a state of active growth to stationary phase. Other abbreviations are listed in FigS4C. B. The graph shows the mean change in the expression status of various lipid classes in WT (blue) and *uip4*Δ (red) by applying log2FoldChange (sample-stationary/sample-log) and p-value<0.05. Error bars represent standard deviation. p-value was used to identify lipids that change significantly with Benjamini and Hochberg multiple testing correction. C. Differential expression analysis was performed in the *uip4*Δ compared to the WT for the lipids identified in the logarithmic phase of growth by applying log2FoldChange. A p-value cut-off of 0.05 was used and was adjusted by Benjamini-Hochberg procedure. The graph shows the mean change in the expression status and error bars represent standard deviation. D. The graph shows the difference between mean expression status of various lipid sub-classes in WT and *uip4*Δ by applying log2FoldChange. The graph shows the mean change in the expression status and error bars represent standard deviation. The abbreviations are defined in the supplementary table. E, F. The bar graphs show the median and internal standard normalized intensity ratio for the identified triacylglycerol (E) and sub-classes of membrane lipids (F) species in the labelled samples. p<0.05*, p<0.005 **, p<0.0005 ***

Differential expression of various lipid classes in the stationary phase versus logarithmic phase was compared between wild type and *uip4*Δ.The hierarchical clustering dendrogram showing the duplicates for three biological replicates of each sample is shown, indicates good technical reproducibility (Fig S4A). The data for lipid species that are unusually high or low is shown in the heatmap (Fig S4B). Several lipid species showed an altered expression in the two growth states and between the two samples (Fig S4B). The abbreviations used are listed in Fig S4C. Intensity ratio averages within each sample group with the top 50 lipids ranked by p-value are shown in Fig S4D. The identified lipids were categorized and expression in the logarithmic phase was used as control. The differences visualized by applying log2 fold change reveals that there were several changes in various lipid classes in both wild type and *uip4*Δ cells, between logarithmic and stationary phase (Fig 6B, S4E). In wild type cells, the synthesis of membrane lipids is reduced and an increase in storage lipids takes place in stationary phase (Fig 6B). However, this regulation is disrupted in the absence of Uip4 (Fig 6B, Fig S4E). When compared to the logarithmic phase of wild type cells, *uip4*Δ has reduced membrane lipids and increased storage lipids such as triacylglycerol (Fig 6C). The lipid class distribution for significantly (p<0.05) altered lipids sub classes (WT-log/*uip4*Δ-log) is shown in Fig 6D. *uip4*Δ cells have high levels of triacylglycerols even in the logarithmic phase (Fig 6E, S5A, S5B). This correlates with the increased number of lipid droplets seen in *uip4*Δ cells. Another major component of lipid droplets are the steryl esters. However, we do not see any major differences in the levels of steryl esters (CE) in the logarithmic phase between wild type and *uip4*Δ (Fig S5C). Most of the membrane lipids such as cardiolipin (CL), phosphatidylglycerol (PG), phosphatidylethanolamine(PE) and phosphatidylcholine (PC) were reduced in *uip4*Δ cells in the logarithmic phase of growth (Fig 6D, 6F). Cardiolipins show a high variability between wild type and *uip4*Δ (Fig S6A,B). While a few of the species have higher concentrations in *uip4*Δ as compared to wild type, on the whole the cardiolipins are downregulated in *uip4*Δ (Fig 6F, S6A, S6B). The cellular sterols also do not show differences between various samples tested (Fig S6C). Only a small, but significant increase in ergosterol peroxide is noticed in *uip4*Δ cells in the stationary phase in comparison to the wild type (Fig S6C). Carnitine is also increased in the *uip4*Δ, particularly in the stationary phase (Fig S6D). Thus, overall it appears that loss of Uip4 results in increased triacylglycerols, generally reduced membrane lipids with minimal effect on sterols and steryl esters.

The changes in lipidome observed could be due to increased synthesis of TAGs or decreased utilization of these or alternately a defect in distributing lipids to organelle membranes. To test if the dysregulation of lipid homeostasis in *uip4*Δ cells is elicited at the genetic level, we performed qRT-PCR analysis of key enzymes involved in various steps of membrane and storage lipid biosynthesis (Fig S7A). However, we did not find any significant difference in the expression of these enzymes in the absence of *UIP4* (Fig S7B, C). There is a discernible downregulation of the two tested triacylglycerols lipase encoding genes, *TGL3* and *TGL4* (Fig S7B,C). It is also possible that the free fatty acid produced are not channelled towards membrane biosynthesis and hence TGs are made to prevent FFA toxicity (Geltinger et al., 2020). This could explain why we have both increased lipid droplets and organelle morphology changes (Fig 3). Therefore, it is likely that Uip4 engages in modulation of lipid homeostasis either directly at the local membrane or indirectly by perturbing organelle contacts otherwise essential for direct transfer of lipid intermediates and enzymes involved in lipid metabolism (Fig 7).

**Figure 7:**
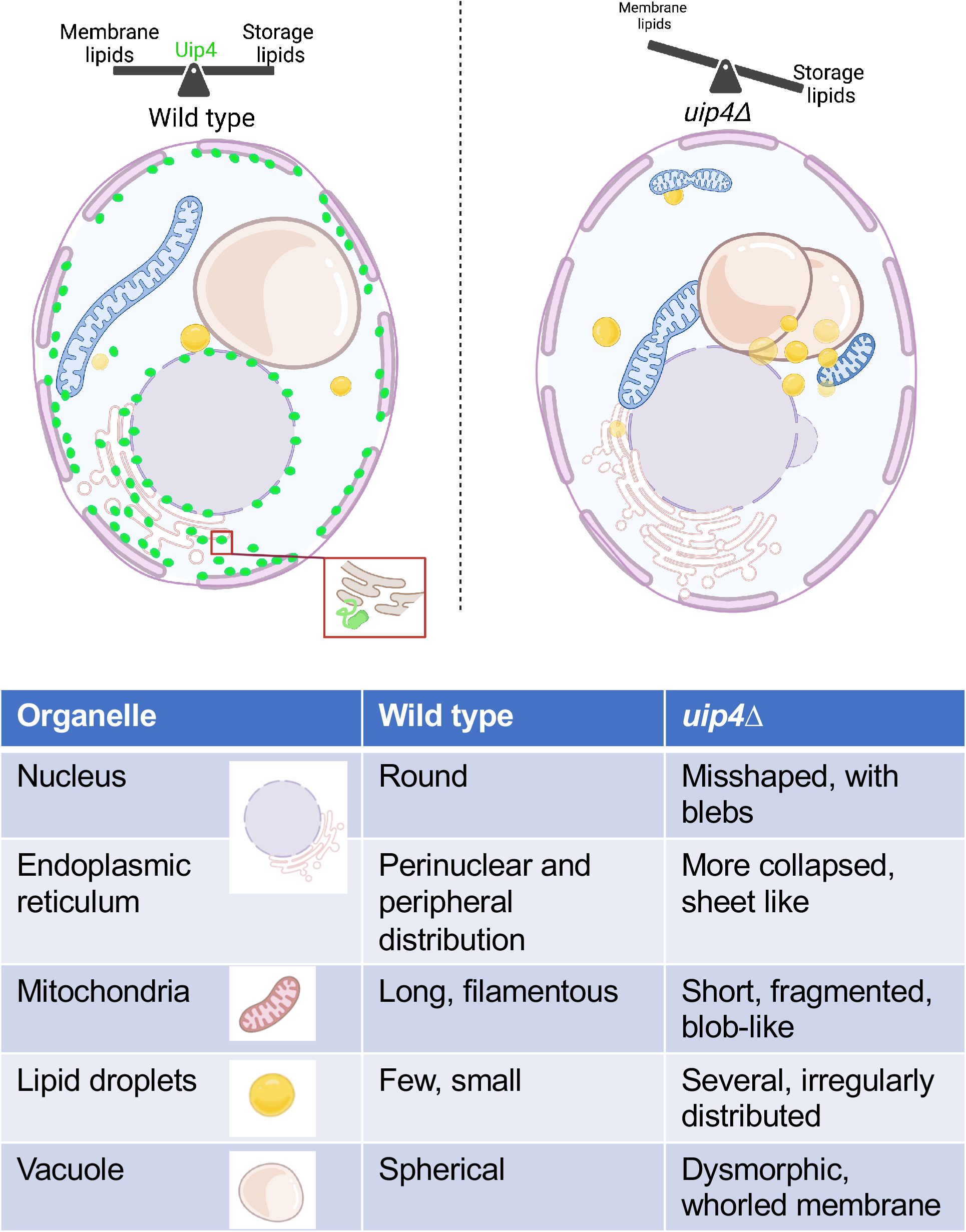
This figure summarises the effects of loss on *UIP4* on organelle shape and contact sites (Created with biorender.com, Agreement number MN256BRXL2). In summary, Uip4 (green), an NE/ER resident protein, is essential for governing the balance between synthesis of membrane lipids and storage lipids in wild type cells (left). This ensures that the organelle morphology and contacts are well established thereby promoting organelle function and cell survival. In the absence of Uip4, the lipid homeostasis is dysregulated, tilting the equilibrium towards increased storage lipid accumulation resulting in abundance of lipid droplets (right). This affects the membrane lipid composition as well, resulting in misshaped and dysfunctional organelles. Unbalanced lipid homeostasis and organelle morphology is concomitant with perturbed membrane contact sites. Our results suggest that Uip4 works as a positive regulator of lipid homeostasis in yeast. The organelles have been marked and the phenotypes are summarised.

## Discussion

In this work, we have identified a growth phase regulated protein, Uip4, in *S. cerevisiae* whose loss leads to reduced survival in stationary phase. Interestingly, loss of this protein that is poorly expressed in logarithmic phase leads to dramatic alterations in multiple organelles even in logarithmic phase of growth. We had earlier demonstrated effects of loss and overexpression of *UIP4* on the nuclear envelope and nuclear pore complexes (Deolal et al., 2022). Now we show that morphology of multiple organelles, including nucleus, lipid droplets, vacuoles, ER and mitochondria is affected (Fig 7). This leads to compromised function of organelles as evidenced by reduced nuclear transport and mitochondrial oxygen consumption rate. In the mutant, abnormally high levels of storage lipids, particularly triacylglycerols, and reduced membrane lipids are seen in the logarithmic phase of growth. We propose that the structural and functional changes seen in multiple organelles in *uip4*Δ are due to this altered lipid homeostasis which results poor post-mitotic survival.

We show that the regulated expression of an ER localized protein, Uip4, is a crucial effector of organelle morphology. Uip4 is expressed more during conditions that limit glucose availability and demand metabolic rewiring. These include reducing glucose concentration in the growth medium, adding galactose or ethanol as carbon source, or cells in stationary phase (Fig 1). Our experimental validation of Uip4 expression during various stages of growth is in line with previous high-throughput reports on significant upregulation of *UIP4* during diauxic shift and stationary phase of growth (Murphy et al., 2015; Di Bartolomeo et al., 2020). We have identified the pathways that regulate expression of Uip4. Interestingly, we find overexpression of Uip4 also leads to dramatic alterations in several organelles (unpublished data), suggesting that the regulated expression of Uip4 is critical for the health of the cell. We find that most of the Uip4 protein colocalizes with ER marker (Fig S3A). However, other studies have detected Uip4 in mitochondria (Reinders et al., 2007; Di Bartolomeo et al., 2020) and lipid droplets (Geltinger et al., 2020). A recent study on identification of novel potential regulators of organelle contact site proteins also found Uip4 as a potential effector of peroxisome-mitochondria and golgi-peroxisome contact site and suggested partial colocalization of a constitutively overexpressed Uip4 tagged at N terminal to these rare and dynamic contact sites (Castro et al., 2022). We did not find any noticeable difference in localization of endogenous Uip4 under altered metabolic states such as logarithmic phase to stationary phase transition at endogenous expression levels (Deolal et al., 2022) and growth on non-fermentable carbon source such as glycerol (data not shown). As this protein in expressed in low levels, establishing precise changes in localization with physiologically relevant changes is challenging. It is likely that Uip4 associates with other organelles in a temporal and dynamic fashion.

Despite our fairly extensive knowledge of basic structural components of organelles, a clear understanding of the mechanisms that contribute towards the maintenance of membrane shape and integrity of associated complexes is lacking even in *S. cerevisiae*. In particular, the dynamic changes in the organelle shape and organisation associated with altered metabolic states are poorly understood. Here we show that altering Uip4 expression is important for maintaining mitochondrial and other organelle morphology during physiologically changing metabolic needs. Particularly, as Uip4 function appears to be important for survival in stationary phase (quiescent stage), it is likely that Uip4 is involved in metabolic reprogramming of cells during this period. Specific mitochondrial proteins like Ups2 have been identified that are expressed when grown in non-fermentable carbon sources and are required for the transfer of phosphatidyl serine to the inner mitochondrial membrane from the outer mitochondrial membrane for phosphatidyl ethanolamine synthesis (Miyata et al., 2016). This suggests there may be multiple such effectors that regulate organelle lipid homeostasis in a metabolic state specific manner. While ER is the major site of cellular lipid synthesis, the biosynthesis of CL, a signature mitochondrial phospholipid, occurs exclusively in the mitochondria (Horvath and Daum, 2013). Altered cardiolipin type and levels observed in *uip4*Δ could be one reason for the altered mitochondrial shape and state as cardiolipin is known to affect both mitochondrial fission and fusion (Ban et al., 2017; Kameoka et al., 2018). The absence of *UIP4* enhances lipid droplets in logarithmic phase of growth (Fig 3C), leads to altered phospholipid composition (Fig 6D) and brings about a reorganisation of organelle contacts (Fig 5). Unlike several proteins that have recently been identified as organelle contacts that contribute to specific lipid transfer (Eisenberg-Bord et al., 2021; Bisinski et al., 2022), we find large changes in lipid droplets and altered lipidome in the absence of *UIP4*. We therefore, speculate that the regulated level of Uip4 is critical to tune the lipid balance between growth and storage. This balance is altered in *uip4*Δ leading to consequences on organelle membrane composition and organelle function which is reflected in the ability of cells to survive under various stress conditions like nutrient depletion and thermal stress.

How does Uip4 bring about all these changes? Uip4 does not have any conserved motifs associated with known enzymatic activity making it challenging to predict the precise molecular function. A prediction of disordered lipid binding regions of Uip4 reveals a high score for its propensity to bind lipids (Katuwawala et al., 2021). It lacks a transmembrane domain and sequence based predictions suggest that the amphipathic region at the C terminal is responsible for membrane association of this negatively charged ER protein and may provide structural stability (Drin et al., 2007; Drin and Antonny, 2010). Additionally, it has a long intrinsically disordered region which might have a role in associating with lipid derivates either directly or in association with other lipid-binding proteins (Deryusheva et al., 2019). Such intrinsically disordered regions of proteins also have short 2-25 amino acid long motifs that can bind other proteins and regulate interactions with other membrane associated proteins. This suggests that Uip4 could interact with both proteins and lipids.

In our attempts to identify interactors of Uip4 using yeast-two hybrid and pull down/MS approaches, we obtained multiple players in protein translation/folding and carbon metabolism. Many of these proteins are also upregulated in stationary phase. Interactors such as Vps13 (involved in protein targeting to vacuole, localizes to organelle membrane contact sites), Vps36 (ESCRT II subunit component), FAA4 (long chain fatty acyl CoA synthase) hint towards involvement of Uip4 in an inter-organellar communication network and potentially lipid transfer. Vps13 is already known to be involved in lipid transfer and regulation of mitochondrial morphology (Bean et al., 2018; Gao and Yang, 2018). It is possible that Uip4 acts a metabolic state specific regulator of functions of Vps13 or other related proteins. Our ongoing studies are aimed at understanding the mechanistic basis of Uip4 function in lipid homeostasis.

Overall, this work raises several important questions. What are the dynamic changes in organelles that are brought about by altered nutrition status? Are new membrane contact sites established? Do existing contact sites interact with different proteins to direct transfers of different metabolites including lipids? It would be imperative to identify proteins that carry out such function but it is a challenging task as they are likely to be expressed transiently and at low levels with potentially redundant functions. Recent large-scale screens to identify contact sites have revealed many new players. More focused screens to identify proteins that are metabolically regulated and their functions in organelle biology would be most useful in unravelling organelle function in metabolic rewiring.

## Materials and Methods

### Yeast growth, strains and plasmids

Yeast strains used in the study are described in Table1, Supplementary file 1. A fresh batch of parent cells were streaked out from the glycerol stock at -80°C before starting an experiment. The cells were grown in either Synthetic Complete (SC) medium containing all amino acids or Yeast Extract-Peptone-Dextrose (YPD) media. Yeast transformation was performed using standard lithium acetate based protocol (Daniel Gietz and Woods, 2002). Strains harbouring plasmids were selected and grown on respective SC plate lacking the selected nutrient. Plasmids used have been described in Table2, Supplementary file 1.

In order to perform growth assays, overnight cultures were sub-cultured to 5mL of fresh medium with equal number of cells. Each of them was allowed to grow in the liquid synthetic complete medium for approximately 4-5 hours at 30℃. The cells were spun and a 10-fold serial dilution was performed. 5µl of cells were spotted on SC plates and incubated for 2-3 days at desired temperature to assess growth differences.

### Protein extraction and western blot

Cells were grown in selection medium either up to mid-log phase or stationary phase, as desired. Proteins were extracted using TCA method of protein precipitation described briefly in (Deolal et al., 2022). Denaturing PAGE was employed to resolve the proteins extracted. Gels for SDS-PAGE were cast using standard protocol. Protein/ epitope tag specific primary and respective secondary antibodies tagged to HRP were used. The signal was detected by chemiluminescence and membrane was imaged in ChemiDoc Imaging system. Densitometric analysis of chemiluminescence readout after western blot was done using Gel Analyzer option of ImageJ/FIJI. Protein expression was normalized to the loading control, usually an antibody to a protein expressed by housekeeping gene. Quantification of the protein expression from western blots of at least 3 independent experiments is performed. Statistical significance was determined by using Student’s t-test to compare the differences between samples, assuming that the population was selected randomly without any bias and data is distributed normally. p-value is reported where necessary and the error bars are representative of the standard error of mean throughout. Uip4 tagged with13xMyc epitope was detected by using α-myc (abcam ab56 (1:5000), ab9106 (1:10000)). Actin (Santa Cruz sc-47778, 1:5000) and tubulin (Abcam ab6160, 1:10000) were used as loading controls.

### qRT PCR

For RNA isolation, a primary culture of desired strains was grown overnight. Next day, appropriate volume of inoculum was taken and secondary culture was set up to an initial OD600 of 0.15-0.20. A 5mL culture containing cells equivalent to 1unit OD600 were harvested after desired growth or treatment. The cell pellet was washed with cold, sterile water, transferred to a microfuge tube and snap frozen in liquid nitrogen. The pellet was then stored in -80°C. To begin RNA isolation, the pellet was resuspended in TES (10mM Tris-Cl pH7.5, 10mM EDTA pH8.0, 0.5% SDS) buffer prepared freshly. This suspension was added to another microfuge tube containing equal volume of water saturated acidic phenol preheated to 65°C. The tubes were then placed in thermomixer set at 65°C and incubated for 1hour with intermittent mixing by gently inverting the tube. The tubes were then placed in ice for 5 min followed by centrifugation at 13000rpm for 5 min. The top aqueous layer was collected and added to an equal volume of chloroform to get rid of residual phenol. The solution was centrifuged again at 13000rpm for 5 min and aqueous layer was collected. To this, 2.5 volumes of chilled 100% ethanol and 0.10 volume of 3M sodium acetate (pH5.2) was added for precipitating RNA. The tubes were then placed in -80°C for at least 2 hours (up to overnight). RNA was pelleted by spinning at 13000rpm for 10 min followed by washing the pellet with 70% ethanol. The pellet was allowed to air dry in the tube followed by resuspension in 100µl of RNase free water. The integrity of RNA was checked by agarose gel. DNase digestion of RNA was done by using RNase free-DNase and absence of DNA contamination was confirmed by PCR. cDNA was prepared by using Verso cDNA synthesis Kit by ThermoScientific using 1µg of RNA as template. The reaction mixture was prepared as per the manufacturer’s protocol. The quality of cDNA was tested by PCR. Transcript levels were tested by qRT-PCR. Primers used are mentioned in Table3, Supplementary file 1. ΔΔCt method was used to calculate fold change (Abraham and Mishra, 2018).

### Glucose estimation

In order to estimate the glycogen and trehalose content, the glucose released upon enzymatic cleavage was measured (Parrou and François, 1997). 10unit OD600 cells were harvested at desired stage of growth. Culture was spun at 3000rpm for 3 min and after removal of culture medium supernatant, the pellet was resuspended in 1ml ice-cold water and transferred to a 2ml microfuge tube. The cells were spun again and the supernatant was removed completely using a vacuum pump. The pellet was resuspended in 250µl of sodium carbonate (0.25M) and incubated at 95°C in a dry bath for 4 hours. Two tubes containing only 250µl sodium carbonate (0.25M), to be used as buffer control were also incubated along with the samples. Next, 150µl of acetic acid (1M) and 600µl of sodium acetate (0.2M) were added to bring the pH to 5.2. This suspension was divided in to four parts, two for enzymatic digestion and 2 for respective ‘no enzyme’ control. 1U/ml of amyloglucosidase from *Aspergilus niger* (Sigma 10115) and 0.025U/ml of trehalase (Sigma T8778) were used to liberate glucose from glycogen and trehalose respectively. The incubation for enzyme digestion was performed overnight at 37°C for trehalase and at 57°C with constant rotation for amyloglucosidase. Next day, the tubes were spun at 13000rpm for 3 minutes. 40µl of suspension was used for glucose estimation. Glucose was measured using Glucose (GO) Assay Kit (Sigma GAGO20) containing glucose oxidase/peroxidase reagent/O-dianisidine reagent in a 96-well plate. Intensity of colour proportional to the original glucose concentration was measured at 540nm.

### Staining organelles with dyes

2unit OD600 equivalent of cells harvested from desired growth conditions were collected by centrifugation and the remaining media was washed off by resuspension in phosphate bufffered saline (PBS), pH7.4. After washing the pellet two times, the cells were resuspended in 500µl of PBS for staining with BODIPY (4,4-difluoro-1,3,5,7,8-pen-tamethyl-4-bora-3a,4a-diaza-s-indacene) 493/503 (final working concentration 2µM), CellTracker Blue CMAC (7-amino-4-chloromethyl-coumarin) (final working concentration 100µM) and mitochondrial dye HC-1 (final working concentration 1µM). The cell suspension was placed in dark with constant gentle rotation, at 30°C for 30 minutes. Then the cells were spun down and washed with PBS thrice to get rid of unbound dye. The pellet was finally resuspended in SC media and images were acquired immediately.

For staining the vacuoles with the vital dye FM4-64 (Invitrogen), the cell pellet was resuspended in YPD containing FM4-64 (final working concentration 8µM) and placed in dark with constant gentle rotation, at 30°C for 30 minutes. Then the cells were spun down and washed with fresh YPD twice and the suspension was again placed in dark with constant gentle mixing, at 30°C for 30-45 minutes. After this 30 minutes of chase, the cells were harvested, washed and resuspended in SC media for imaging.

### Microscopy

For live cell imaging, cells resuspended in SC media were mounted onto a 35mm cover glass bottom dish coated with 0.1% concanavalin A solution (Deolal and Mishra, 2022). After allowing the cells to settle, unbound cells were rinsed with SC media. Images were acquired in confocal microscope equipped with a temperature-controlled stage set at 30°C. All images were acquired on Leica TCS SP8 using HC PL APO CS2 63X/1.40 OIL objective. Optimal laser power, image size and acquisition settings were used. At least three biological replicates for each sample were imaged for quantification and analysis. Deconvolution was performed using Huygens Professional version 17.04 (Scientific Volume Imaging, The Netherlands, http://svi.nl) (Deolal and Mishra, 2022). For transmission electron microscopy (TEM), yeast cell cultures were grown to mid-log phase 0.5 OD600. Samples were fixed as described in (Deolal et al., 2022). The sectioning and image acquisition was carried out at TEM facility of ACTREC, Tata Memorial Centre, Mumbai.

### Image quantification and analysis

3 biological replicates were imaged for each condition and panels representative of the phenotype are cropped and shown. Number of lipid droplets were counted using the Find Maxima function of FIJI. The diameter of single cells was measured using the Straight line tool. For the intensity profile, the data was extracted using the Plot Profile function.

### Lipid estimation and analysis

#### Sample preparation

Wild type and *uip4*Δ cells were grown to either mid-log (OD600-0.6) or stationary phase (48 hours grown culture) in SC medium. 3unit OD600 equivalent cells were harvested in a 2mL microfuge tube by centrifuging the tubes at 6000rpm for 10 minutes at 4°C. The media was removed and cells were resuspended in 1.5mL of cold HPLC grade water. The tubes were again centrifuged at 6000rpm for 10 minutes at 4°C and this washing was done 3 times. After final wash, the tubes containing cell pellets were placed in liquid nitrogen for two minutes to ensure that they are metabolically inactive. Then the cell pellets were stored in -80°C freezer until analysis. Simultaneously, three blank samples were prepared by washing empty microfuge tubes in a manner similar to those containing cell pellets. All tubes were protected from light. The samples were shipped to The Metabolomic Innovation Centre (TMIC), Canada for further lipid extraction and identification.

### Lipid Extraction

The extraction was performed strictly following the protocol based on a modified Folch liquid-liquid extraction protocol. Each aliquot of 0.57 to 1.82 mg of sample was mixed with NovaMT LipidRep Internal Standard Basic Mix for Tissue/Cells (an internal standard mixture composed of 15 deuterated lipids), dichloromethane and methanol. A clean-up step was performed with water. Samples were equilibrated at room temperature for 10 min and centrifuged at 16,000 g for 10 min at 4°C. An aliquot of the organic layer was evaporated to dryness with a nitrogen blowdown evaporator. The residue was immediately re-suspended in NovaMT MixB, vortexed for 1 min, and diluted with NovaMT MixA.

A pooled mixture composed by one aliquot of the organic extract from each sample was prepared for quality control (QC). The pooled mixture was split into equal aliquots, evaporated to dryness with a nitrogen blowdown evaporator, purged with nitrogen and stored at -80°C. One QC aliquot was resuspended with each randomized batch of six or nine samples (4 batches). The six or nine samples within each batch were injected in between two injection replicates of the corresponding QC aliquot. Multiple QC aliquots were also injected before and after all samples to ensure technical stability.

### LC-MS analysis

The LC-MS analyses were performed by strictly following the SOP in both positive and negative ionization. Sample extracts were injected between injection replicates of the QC pooled mixture prepared with the same sample batch. A total of 30 sample injections (experimental duplicates of 15 samples) and 14 QC injections (7 aliquots of the pooled mixture injected in duplicates) were performed in each ionization polarity. MS/MS spectra were acquired for all samples for identification. Lipid features were extracted and aligned using NovaMT LipidScreener. A three-tier identification approach based on MS/MS spectral similarity, retention time and accurate mass match was employed for lipid identification. A nine-tier filtering and scoring approach embedded in NovaMT LipidScreener was employed to restrict the number of matches and select the best identification. A three-tier identification approach based on MS/MS spectral similarity, retention time and accurate mass match was employed for lipid identification. Molecules in Tier 1 were identified by MS/MS match with score ≥500 and precursor mass error ≤20.0 ppm and 5.0 mDa. Molecules in Tier 2 were identified MS/MS match with score <500 and precursor mass error ≤20.0 ppm and 5.0 mDa. Lipids in Tier 3 were identified by mass match with m/z error ≤20.0 ppm and 5.0 mDa. Features identified in tier 1 and 2 were used for all the analysis described here (data provided as supplementary excel 2). Quality control check was done by plotting Principal Component Analysis scores and hierarchical clustering dendrogram.

### Data Normalization

Identified features were normalized by internal standards and the median intensity ratio. First, data normalization of identified features was performed by using a set of 15 deuterated internal standards belonging to different lipid classes (NovaMT LipidRep Internal Standard Basic Mix for Tissue/Cells). The positively and putatively identified lipids were matched to one of the 15 internal standards according to lipid class similarity and expected retention time range for each class. Intensity ratios, i.e., intensity of each lipid divided by intensity of the matched internal standard, were calculated for internal standard normalization. Second, the identified features were median-normalized, i.e., the intensity ratios for each identified feature were divided by the median intensity ratio of each sample experiment. Statistical analysis was performed with MetaboAnalyst 5.0 (https://www.metaboanalyst.ca/) and LipidSig (http://chenglab.cmu.edu.tw/lipidsig/). Hierarchical clustering dendrogram, heat map and volcano plots shown in Fig S4 were downloaded from MetaboAnalyst. Graphs were plotted in GraphPad Prism 8.0.

### Metabolic flux analysis using Agilent Seahorse

The oxygen consumption rate (OCR) and extracellular acidification rate (ECAR) of wild type and *uip4*Δ cells was measured using the Agilent XFp metabolic flux analyser. The assay involved injection of glucose (20%) or lactate (20%) using the built-in injection ports on XFp sensor cartridges. The assay was performed as per the Agilent Seahorse XFp User Guide with certain modifications suited for yeast system. Filter sterilized synthetic complete medium (pH ∼7.4) containing 2% glucose was used as the assay medium. Equal number of cells were used for WT and *uip4*Δ. The OCR data generated by the Wave software was exported in Prism format and plots were generated.

## Supporting information

List of Strains, plasmid and primer

## Acknowledgement

Authors would like to thank Dr. Liang Li and the staff at The Metabolomic Innovation Centre, University of Alberta, Canada for Lipidomics of yeast cells at different phases of growth (Project TMIC01×8) and Dr. Sharada Sawant, Electron Microscopy Facility (Advanced Centre for Treatment, Research & Education in Cancer, Tata Memorial Centre, Mumbai) for TEM support . We would like to acknowledge Diane McVey Ward-University of Utah and Maya Schuldiner-Weizmann Institute of Science for strains.

## Author contributions

PD contributed to the design of study, acquisition, analysis and interpretation of data. KR performed all the qRT-PCR experiments. BD generated western blots shown in S1(E-G). KM contributed to the conception and design, analysis and interpretation of data, and acquisition of funding. PD and KM wrote the manuscript. All authors read and approved the final manuscript.

## Declaration

The authors declare that they have no competing interests.

---

**Figure S1:**
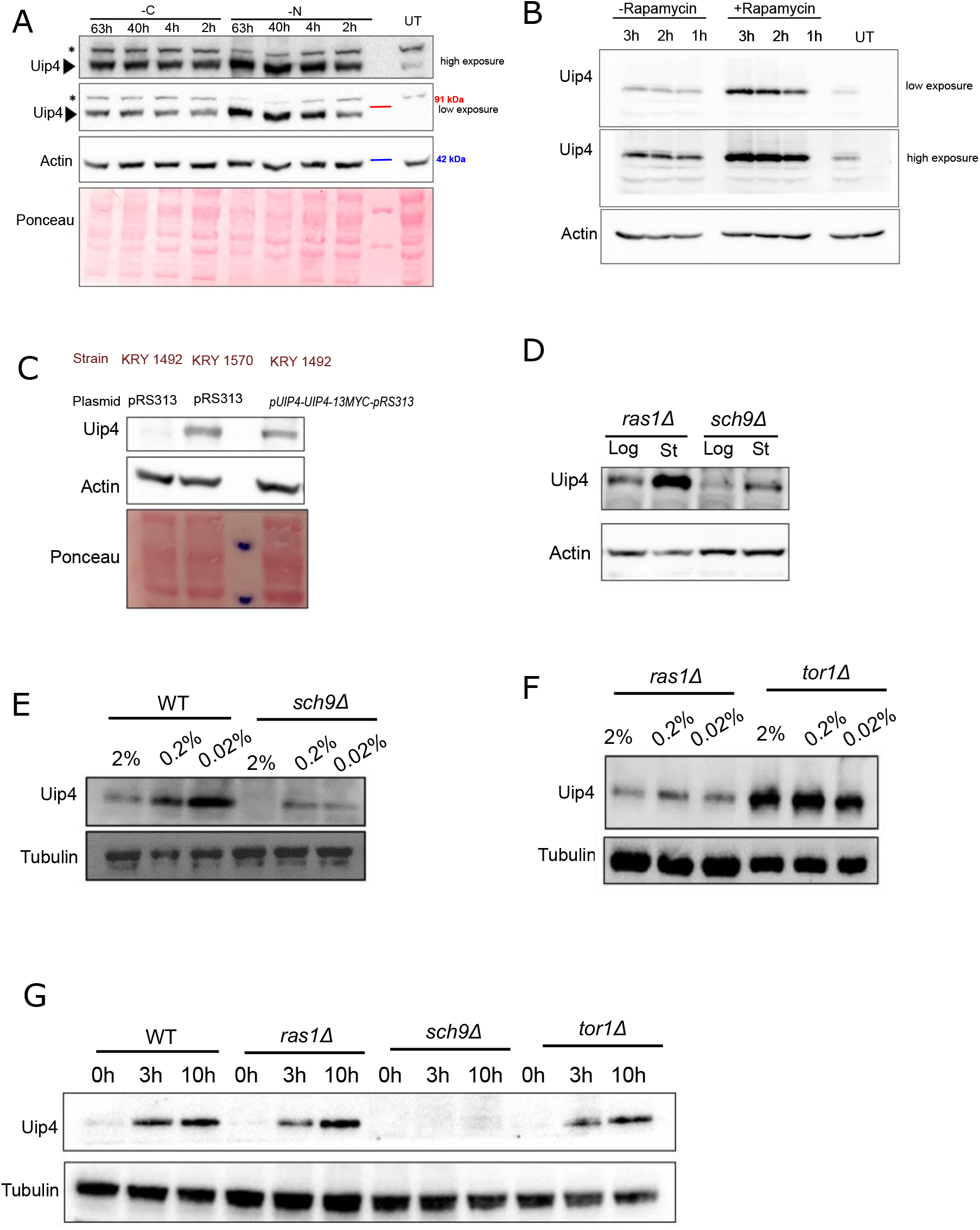
A. Uip4-13Myc tag bearing cells were harvested from mid-log phase (UT) and transferred to a media with either limiting carbon (C) or nitrogen (N) source. Western blot showing the expression levels of Uip4 in the cells harvested at the indicated time points is shown. The non-specific band is indicated by *. Band corresponding to Uip4-13Myc is marked by an arrow. Ponceau staining shows the total protein in the extract. B. Uip4-13Myc tag bearing cells were harvested from mid-log phase (UT) and transferred to a media with either 200ng/ml of Rapamycin (dissolved in DMSO) or equivalent volume of DMSO. Western blot showing the expression levels of Uip4 in the cells harvested at the indicated time points is shown. C. The western blot shows expression of Uip4 from its native promoter either at the genomic locus (lane 2) or from a centromeric plasmid (lane 4). Wild type cells transformed with empty vector serves as a negative control (lane 1). Lane 3 has molecular weight marker. Ponceau staining shows the total protein in the extract. D. Cells from *ras1*Δ and *sch9*Δ strains were harvested either during logarithmic phase or stationary phase of growth as indicated. Western blot showing the expression levels of Uip4 in the cells harvested at the indicated growth points is shown. E, F. Indicated strains expressing Uip4-13xMyc from its endogenous loci were grown to mid-log phase in 2% glucose and then shifted to media containing varying concentrations of glucose (2%, 0.2%, 0.02%) Cells were harvested after 3 hours and Uip4 levels were checked by western blot. α-myc was used to detect Uip4 and α-tubulin was used as a loading control for total protein. G. . Indicated strains expressing Uip4-13xMyc from its endogenous loci were grown to mid-log phase in 2% glucose and then shifted to media staved for nitrogen (0.17% YNB, 2% glucose) Cells were harvested after 3 hours and 10 hours, and Uip4 levels were checked by western blot. α-myc was used to detect Uip4 and α-tubulin was used as a loading control for total protein.

**Figure S2:**
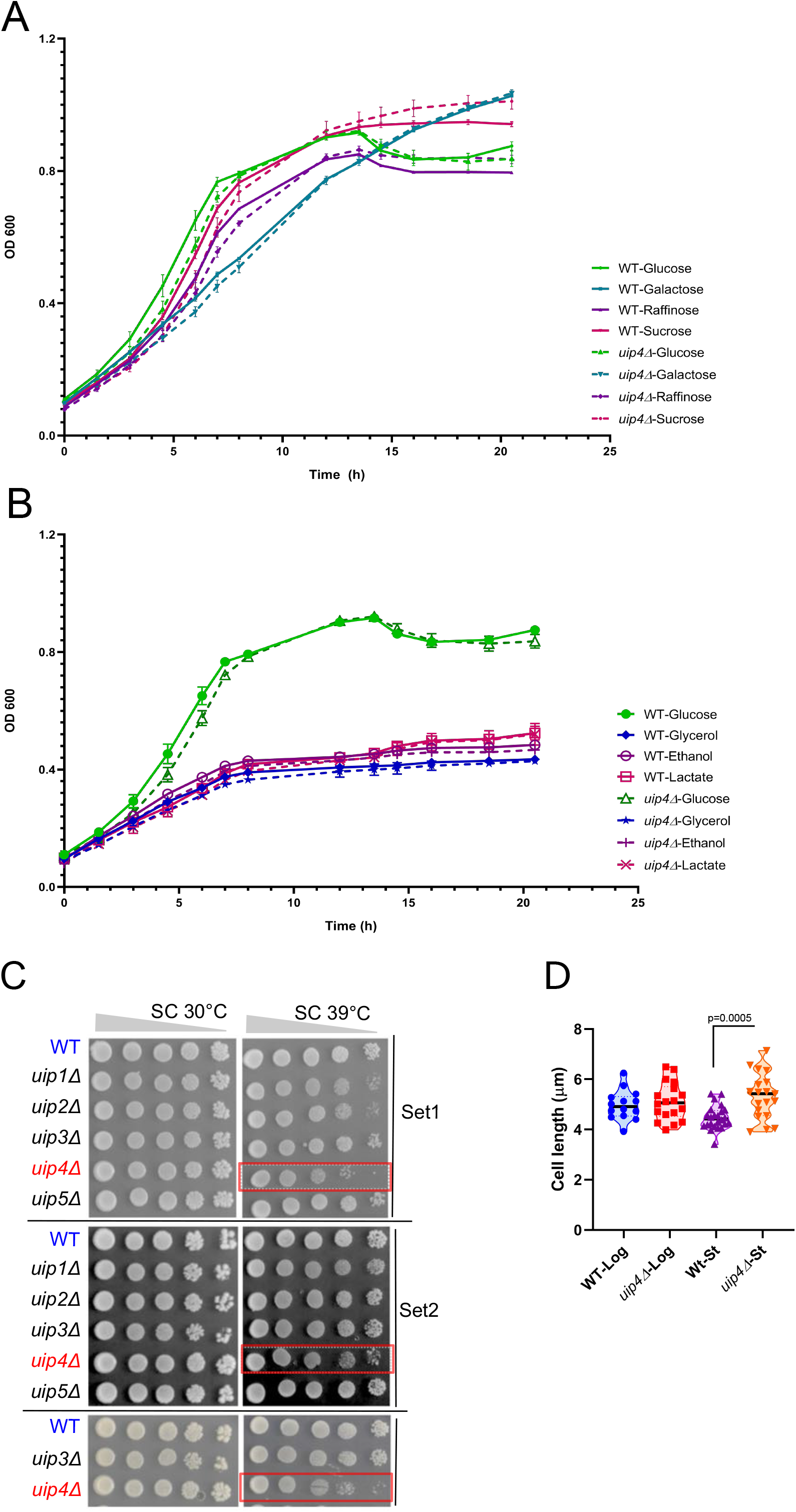
A, B. Growth of wild type and *uip4*Δ cells was monitored in media containing various fermentable (A) and non-fermentable (B) carbon sources and compared to the standard 2% glucose containing medium. OD600 is plotted with growth time. C. Serially diluted cultures of the indicated strains were taken and 5µl of 10-fold serial dilutions were spotted on a SC-plate. The plates were incubated at either 30°C for 2 days or 39°C for 3 days. Three sets of plates showing growth of serially diluted cultures is shown. *uip4*Δ highlighted to indicate the lowered growth at 39°C. D. A plot showing the distribution of cell length is shown. Diameter of wild type and *uip4*Δ cells during logarithmic and stationary phase are compared. Horizontal line indicates mean.

**Figure S3:**
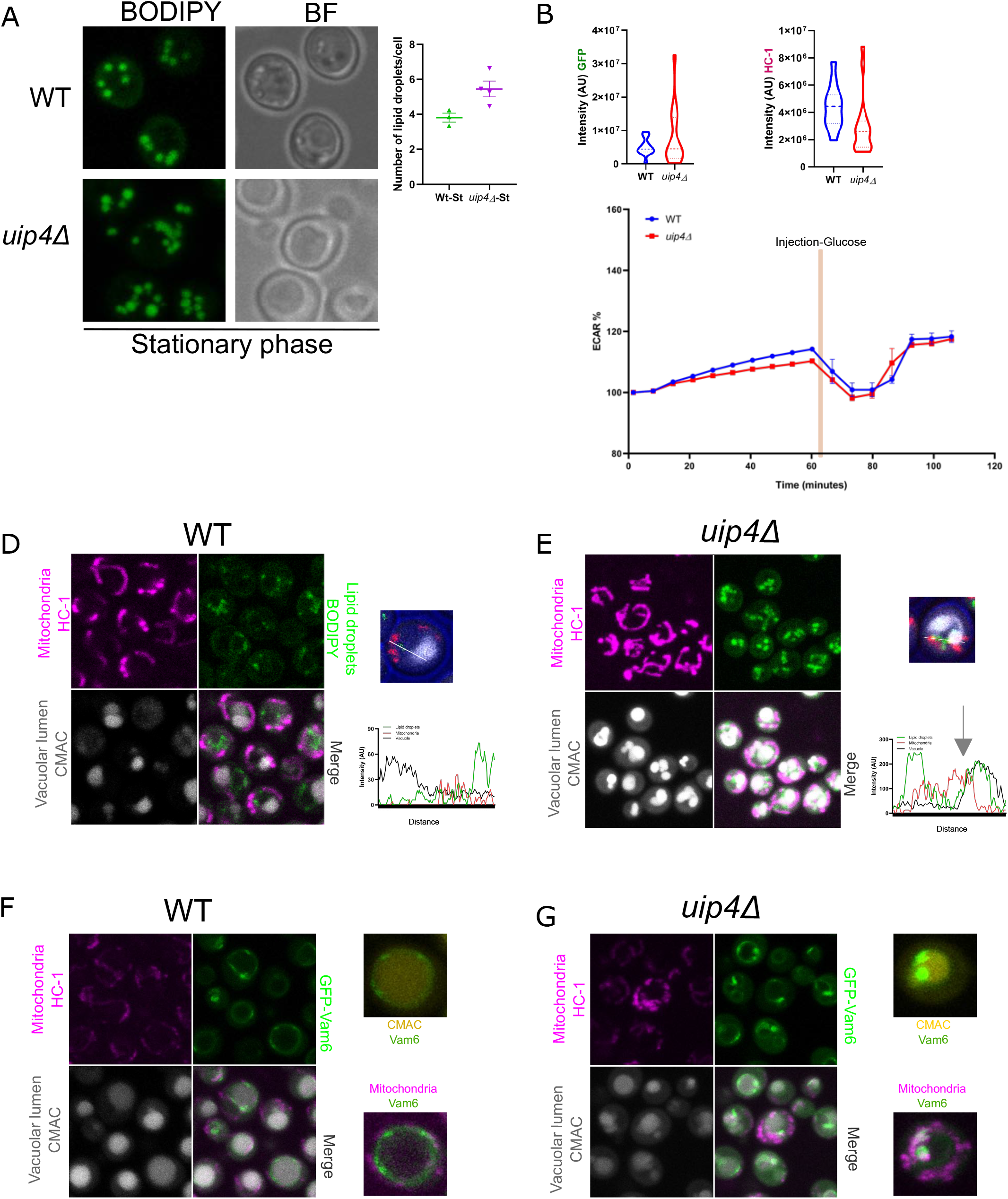
A. Indirect immunofluorescence was performed to visualize localization of Uip4, in cells bearing *UIP4-13MYC* at the endogenous loci, grown in medium containing 2% glucose. α-myc is used to detect Uip4 and α -Pdi1 is used as a marker for ER. Scale bar-1µm. B. Lipid droplets in WT and *uip4*Δ harvested at stationary phase of growth were stained using BODIPY 493/503. Maximum intensity projections of representative images are shown. Scale-2µm. A plot showing distribution of number of lipid droplets per cell is shown on the right. Horizontal line indicates mean. C. Integrated mean fluorescence densities of Su.9-Mito-GFP and HC-1 staining for WT and *uip4*Δ cells shown in Fig 3E is plotted. n=30-40 cells. AU-arbitrary units. D. The glycolytic function of wild type and *uip4*Δ cells was tested by directly measuring the extracellular acidification rate (ECAR) of the cells on the Seahorse XFp Analyzer. The ECAR profile normalized with values at the beginning of the workflow as baseline are plotted. The injection is an addition of glucose (111mM) in the medium. E, F. WT and *uip4*Δ cells harvested during logarithmic phase of growth were co-stained with HC-1, BODIPY and CMAC-blue to label mitochondria, lipid droplets and vacuole lumen respectively. The relative distribution of the three signals as seen in majority of the cells for each strain in shown as the line profile plot of fluorescence intensity across the distance of the line drawn in the cell shown on right. G, H. WT and *uip4*Δ cells expressing GFP-Vam6 harvested during logarithmic phase of growth were co-stained with HC-1 and CMAC-blue to label mitochondria and vacuole lumen respectively. To insets show co-staining of Vam6 (green) with either Vacuole lumen (yellow) or mitochondria (magenta).

**Figure S4:**
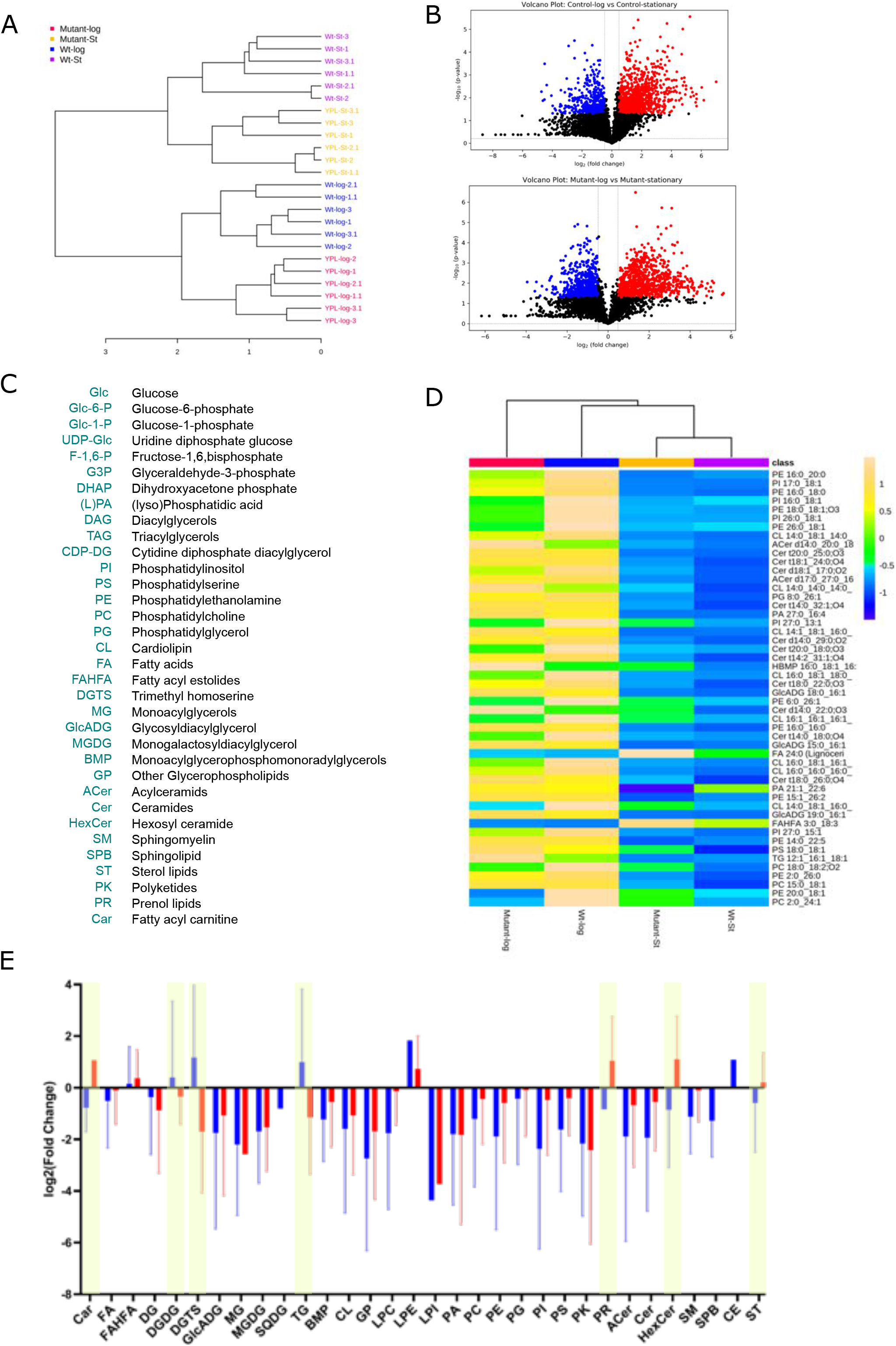
A. The hierarchical clustering dendrogram showing the duplicates for three biological replicates of each sample is shown. The clustering indicates good technical reproducibility. B. The volcano plots shown was generated by plotting the fold change of each lipid, calculated as Mean (Wt-log)/Mean (Wt-stationary) against the p-value for each lipid (top). When using FC (Control-log / Control-stationary) >1.40 or <0.71, p-value <0.05, and p-value adjusted for false-discovery rate (q-value) <0.25 as the criteria for significance, the analysis resulted in 418 significantly altered lipids with FC <0.71 (blue); and 931 significantly altered lipids with FC >1.40 (red). The p-value threshold of 0.05 corresponded to a minimum q-value of 0.07. The volcano plots shown were generated by plotting the fold change of each lipid, calculated as Mean (*uip4*Δ-log)/Mean (*uip4*Δ-stationary) against the p-value for each lipid (bottom). When using FC(Mutant-log / Mutant-stationary) >1.40 or <0.71, p-value <0.05, and q-value <0.25 as the criteria for significance, the analysis resulted in 855 significantly altered lipids with FC <0.71 (blue); and 876 significantly altered lipids with FC >1.40 (red). The p-value threshold of 0.05 corresponded to a maximum q-value of 0.04. C. The metabolites mentioned in Fig 6A and abbreviations for various lipid species are described. D. Heatmap for intensity ratio averages of each group is shown for all samples with the top 50 lipids ranked by p-value. The scale is shown on the right and the samples are labelled at the bottom of the heat map. Each coloured cell corresponds to a concentration value in the mean centred data with samples in rows and lipid species in the columns. Intensity ratio averages within each sample group with the top 50 lipids ranked by p-value are shown. The significance of the change was determined by using t-test/ANOVA. E. The graph shows the mean change in the expression status of various lipid sub-classes in WT (blue) and *uip4*Δ (red) by applying log2FoldChange (sample-stationary/sample-log) and p-value<0.05. Error bars represent standard deviation. p-value was used to identify significant lipids with Benjamini and Hochberg multiple testing correction. The sub-classes showing contrasting expression from logarithmic phase to stationary phase between the two samples are highlighted in yellow.

**Figure S5:**
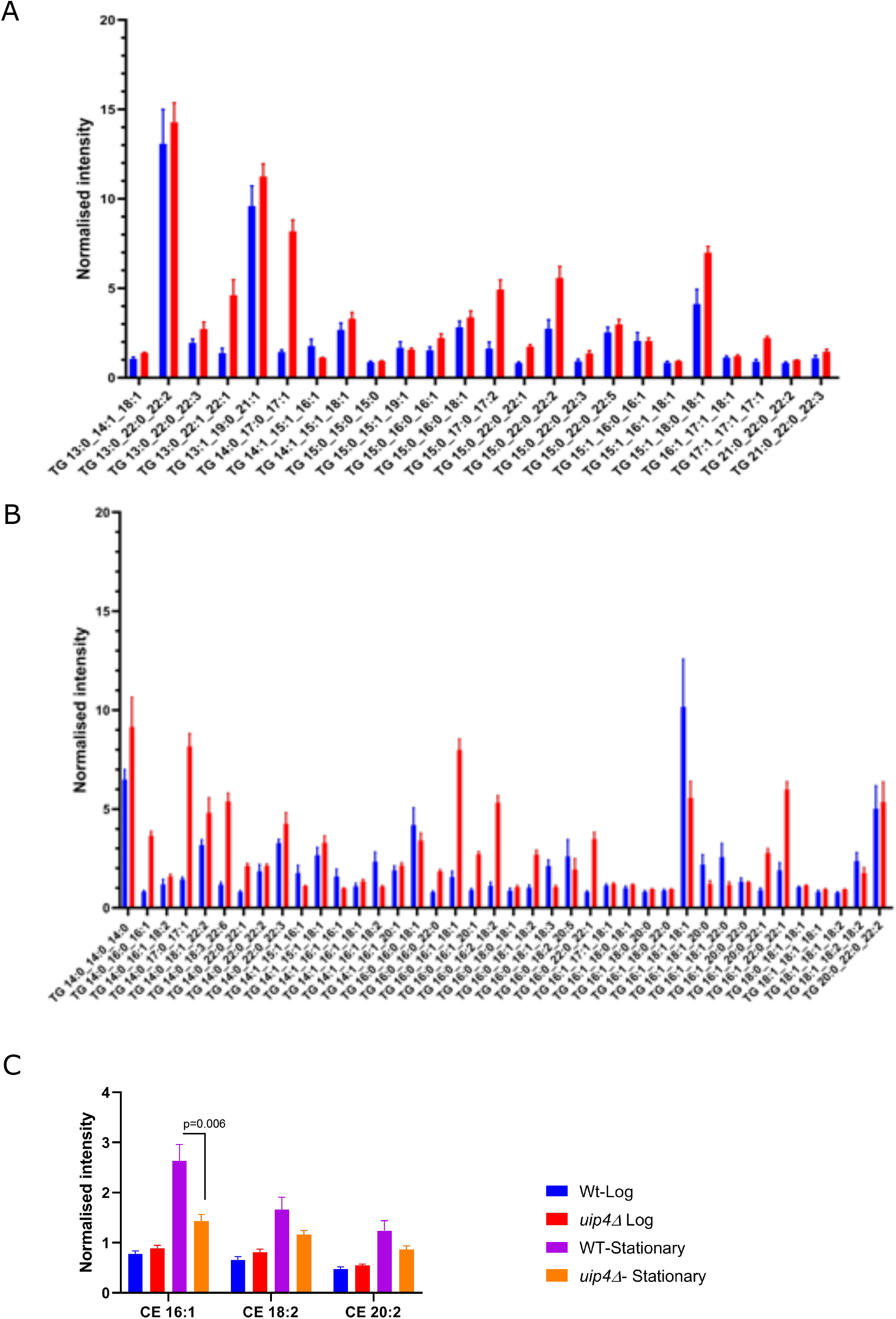
A-C. The bar graphs show the median and internal standard normalized intensity ratio for the identified lipid species in the labelled samples. TG-triacylglycerols, CE-steryl esters.

**Figure S6:**
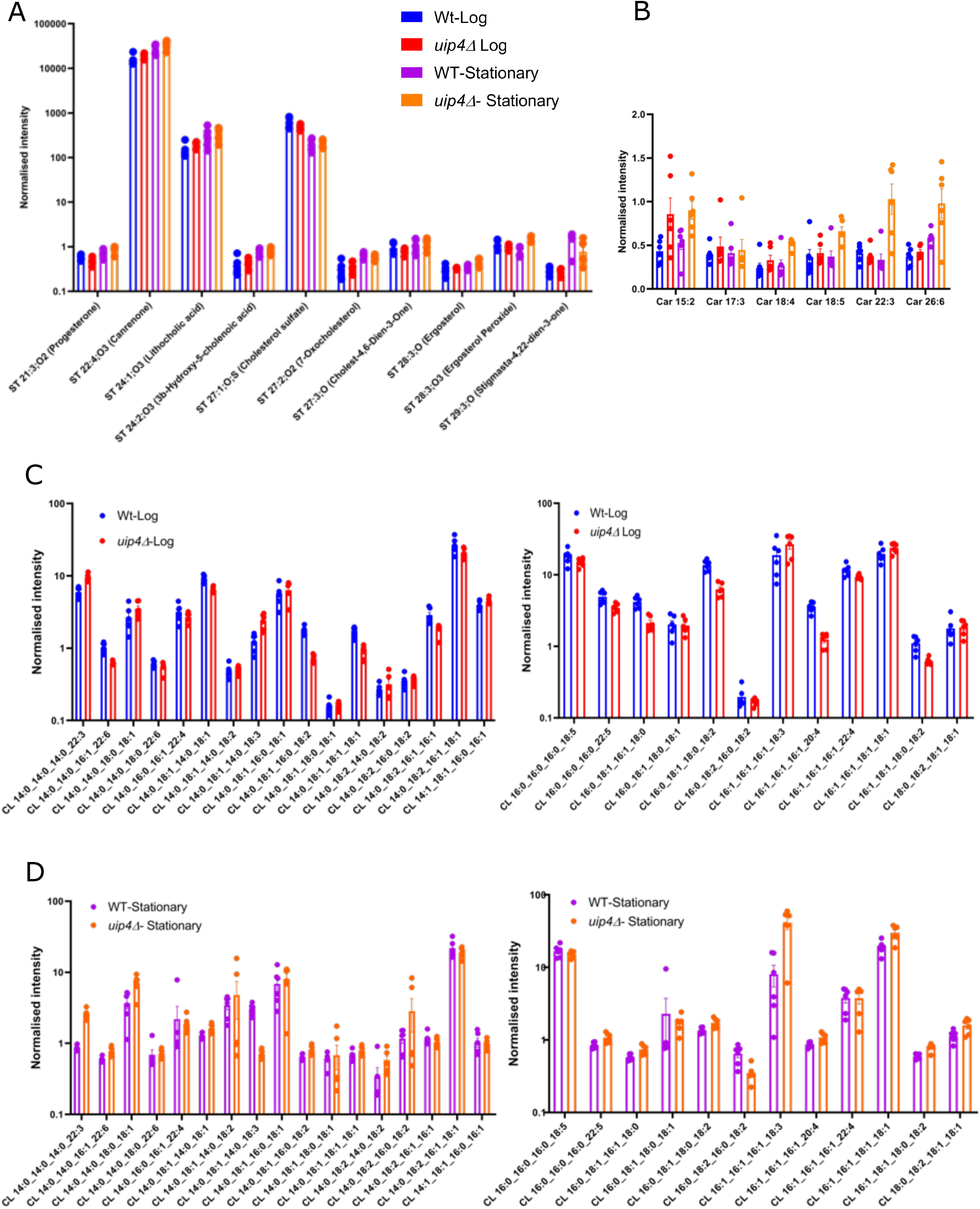
A-D. The bar graphs show the median and internal standard normalized intensity ratio for the identified lipid species in the labelled samples. CL-cardiolipin, ST-sterol lipids, Car-fatty acyl carnitine.

**Figure S7:**
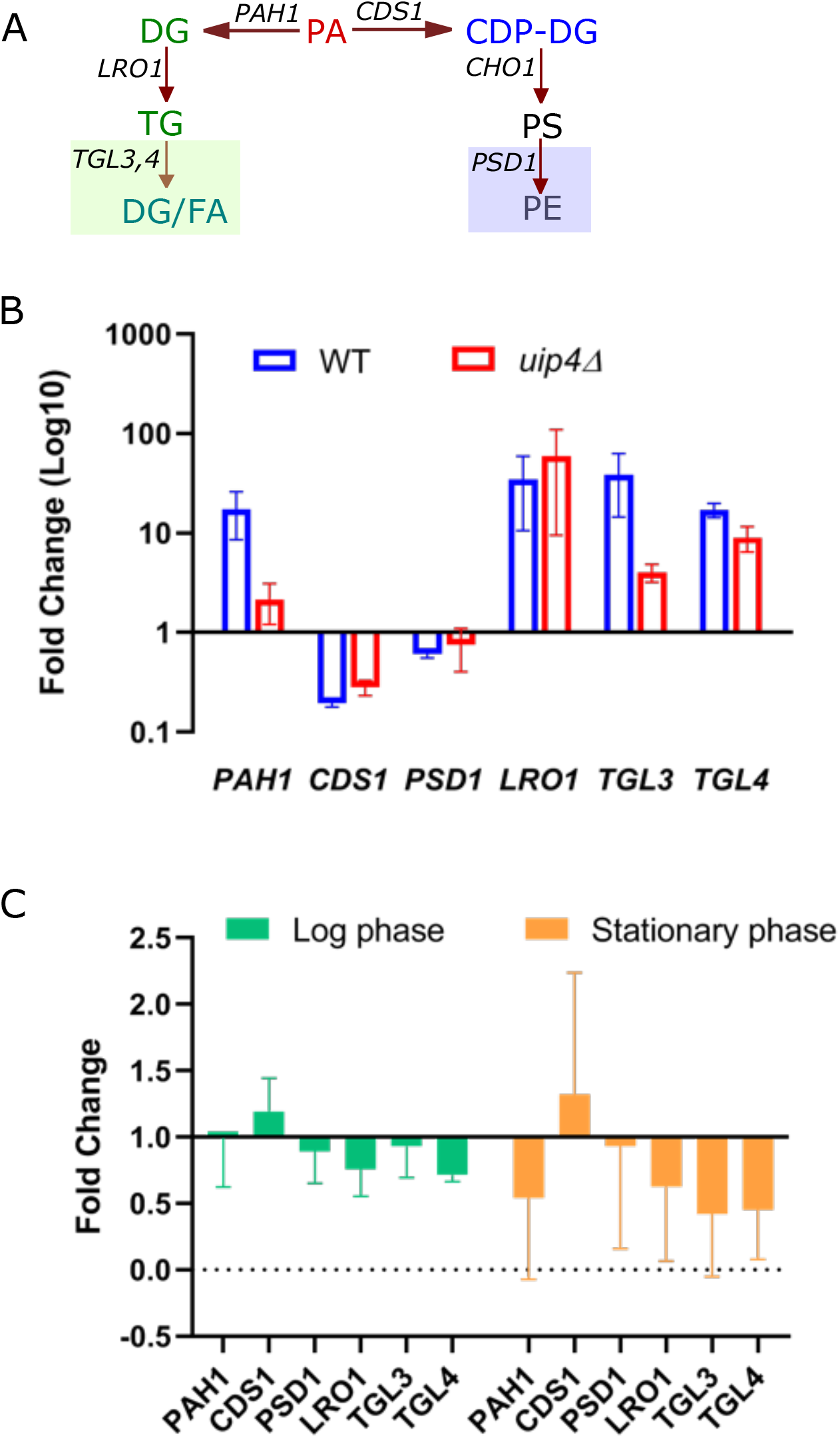
A. A schematic representation of some of the enzymes involved in biosynthesis of storage (green) and membrane lipids (blue) from the precursor phosphatidic acid (red) is shown. B, C. RNA was isolated from WT and *uip4*Δ strains at mid-log phase and stationary phase (48hours) and qRT-PCR was performed using cDNA. *ACT1* was used as normalizing control. The histogram in B shows abundance of indicated transcripts in stationary phase relative to mid-log phase. The comparison of abundance of transcripts in *uip4*Δ relative to wild type in the two growth phases is shown in C. The differences between the data sets are not statistically significant (p>0.05).

